# Spotiflow: accurate and efficient spot detection for fluorescence microscopy with deep stereographic flow regression

**DOI:** 10.1101/2024.02.01.578426

**Authors:** Albert Dominguez Mantes, Antonio Herrera, Irina Khven, Anjalie Schlaeppi, Eftychia Kyriacou, Georgios Tsissios, Evangelia Skoufa, Luca Santangeli, Elena Buglakova, Emine Berna Durmus, Suliana Manley, Anna Kreshuk, Detlev Arendt, Can Aztekin, Joachim Lingner, Gioele La Manno, Martin Weigert

**Affiliations:** Institute of Bioengineering, School of Life Sciences, École Polytechnique Fédérale de Lausanne (EPFL), Lausanne, Switzerland; Brain Mind Institute, School of Life Sciences, École Polytechnique Fédérale de Lausanne (EPFL), Lausanne, Switzerland; Bioimaging and Optics Platform, École Polytechnique Fédérale de Lausanne (EPFL), Lausanne, Switzerland; Swiss Institute for Experimental Cancer Research, School of Life Sciences, École Polytechnique Fédérale de Lausanne (EPFL), Lausanne, Switzerland; Developmental Biology Unit, European Molecular Biology Laboratory (EMBL), Heidelberg, Germany; Cell Biology and Biophysics Unit, European Molecular Biology Laboratory (EMBL), Heidelberg, Germany; Institute of Physics, School of Basic Sciences, École Polytechnique Fédérale de Lausanne (EPFL), Lausanne, Switzerland; Center for Organismal Studies (COS), University of Heidelberg, Heidelberg, Germany; Center for Scalable Data Analytics and Artificial Intelligence (ScaDSAI), TU Dresden, Dresden, Germany

**Author notes:** Correspondence to:, M.W. ( / @martweig; G.L.M. ( / @GioeleLaManno / gioelelamanno.com).

## Abstract

Identifying spot-like structures in large and noisy microscopy images is a crucial step to produce high quality results in various life-science applications. Imaging-based spatial transcriptomics (iST) methods, in particular, critically depend on the precise detection of millions of transcripts in images with low signal-to-noise ratio. Despite advances in computer vision that have revolutionized many biological imaging tasks, currently adopted spot detection techniques are mostly still based on classical signal processing methods that often lack robustness to changing imaging conditions and thus require tedious manual tuning per dataset. In this work, we introduce Spotiflow, a deep learning method that achieves subpixel-accurate localizations by formulating the spot detection task as a multi-scale heatmap and stereographic flow regression problem. Spotiflow can be used for 2D images and 3D volumetric stacks and can be trained to generalize across different imaging conditions, tissue types and chemical preparations, while being substantially more time- and memory-efficient than existing methods. We show the efficacy of Spotiflow via extensive quantitative experiments on a variety of diverse datasets and demonstrate that the enhanced accuracy of Spotiflow leads to meaningful improvements in the biological insights obtained from iST and live imaging experiments. Spotiflow is available as an easy-to-use Python library as well as a napari plugin at https://github.com/weigertlab/spotiflow.

## Introduction

Many methods in the life sciences produce images for which detecting and localizing spot-like objects is a crucial step preceding specialized downstream analyses. This problem, commonly referred to as *spot detection* [1–4], has been the computational basis of many methods in genomics over the last decades [5, 6]. Recently, the advent of imaging-based spatial transcriptomics (iST) has brought this task to a significantly more challenging and computationally demanding domain [7] (Fig. 1a). In iST, RNA molecules are located in situ within large tissue sections during sequential imaging cycles to generate gene expression maps at subcellular resolution [8–10]. Popular iST techniques such as MERFISH [8], seqFISH [10] or HybISS [9] require the detection of millions of spots in gigabyte-sized images with high accuracy, sensitivity, and computational efficiency. Due to the preservation of the native tissue context, any general iST spot detection method has to address multiple imaging challenges such as autofluorescence background, aspecific probe binding, or inhomogeneous spot sizes and densities (Supp. Video 1). High sensitivity and accuracy are crucial because, for most iST methods, transcript identity is encoded combinatorially in the sequence of multiple multi-channel images across different imaging rounds [8, 9]. As a result, suboptimal spot detection performance in one channel or imaging round can significantly reduce sensitivity and lead to transcript identity misattribution [11]. Furthermore, the advent of 3D-capable technologies such as STARmap [12] or EASI-FISH [13] yields specific challenges such as voxel anisotropy, asymmetrical axial profiles of the point-spread function or depth-depending illumination.

**Figure 1:**
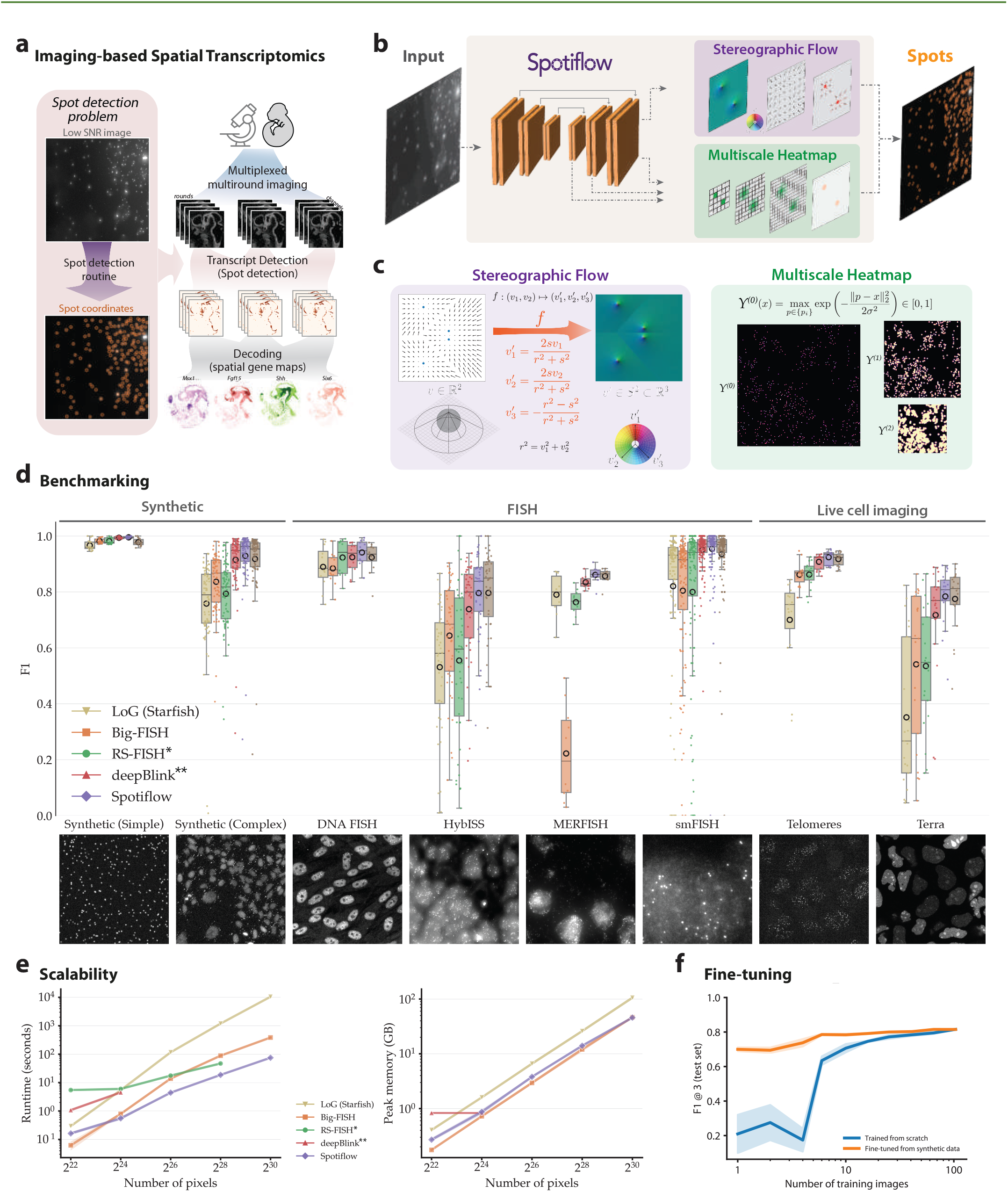
Fast, scalable, and accurate fluorescence spot detection with Spotiflow. **a)** Depiction of a common processing pipeline for imaging-based spatial transcriptomics (iST) data, in which spot detection is a critical step. **b)** Spotiflow is trained to detect spots from microscopy images via two different synergic tasks, *multiscale heatmap regression* and *stereographic flow regressions)*. **c)** The ground truth objects to be regressed are computed from point annotations {*p*_*k*_}. First, a full-resolution Gaussian heatmap *Y*^(0)^ is obtained by generating isotropic Gaussian distributions of variance *σ*^*2*^ centered at spot locations. This Gaussian heatmap is further processed to obtain lower resolution versions, yielding *multiscale heatmaps Y*^(*l*)^, which are all regressed. Second, a local vector field V = {υ_*ij*_} is built by placing a vector directed to the closest spot center at every pixel of the image. We obtain the *stereographic* 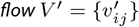 by computing, position-wise, the inverse stereographic projection *f* of the local vector field. **d)** Benchmarking of spot detection methods on different datasets, grouped by their modality (*Synthetic, FISH, Live cell imaging*). Shown is the distribution of F_1_ scores per image in the test set of every dataset (higher is better, *cf*. Supp. Note 3.2). A sample training image is depicted under each dataset. Each method was optimized/trained and tested individually in each dataset except Spotiflow (*general*), which was trained on all datasets except *DNA FISH*. For the *DNA FISH* dataset, the reported Spotiflow *general* score on the test set was obtained with the *general* model fine-tuned on three training images (∼14% of the whole train set). **e)** Runtime (top) and peak CPU memory (bottom) assessment for different methods at different image sizes. Parameters of each method were calibrated so that the amount of detections were in the same order of magnitude. deepBlink and Spotiflow were profiled on GPU, the other methods on CPU. Memory usage excludes constants such as library imports or interpreter initialization. ^*^ RS-FISH not shown for sizes >32k due to Java size-related limitations. RS-FISH memory was not profiled as the implementation is not in Python. ^**^ deepBlink could not be run for sizes > 4k due to GPU memory limitations. **f)** F_1_ score on live-cell dataset *Terra* after fine-tuning a Spotiflow model pre-trained on *synthetic-complex* with an incrementally increasing number of out-domain training images from *Terra*.

Commonly used spot detection pipelines for iST often rely on classical threshold-based methods such as Laplacian-of-Gaussian (LoG) [14, 15] or radial symmetry [16]. While these approaches perform well on simulated or relatively clean data, they often struggle with realistic images that exhibit artifacts, autofluorescence, and varying contrast. Although a few deep learning-based methods have been proposed for this task [17–20], they are often hard to use, do not provide subpixel accuracy (with the notable exception of [19]), and don’t scale to gigapixel-sized images. Moreover, no existing learning-based method allows end-to-end modelling of 3D volumes, with most of the methods resorting to a suboptimal Z-blending strategy (*e.g*. [19]). Overall, the classical iST spot detection methods currently used lack robustness in challenging image conditions, are computationally inefficient for large images, and require manual threshold tuning for every channel and imaging round, which limits their applicability and accuracy in large-scale iST experiments.

Here we introduce Spotiflow, a deep learning-based, threshold-agnostic, and subpixel-accurate spot detection method that outperforms other commonly used methods on a variety of iST and non-iST modalities while being substantially more time and memory efficient than other solutions, especially in the large size regime. Spotiflow is trained to predict *multi-scale Gaussian heatmaps* and exploits a novel *stereographic flow* regression task from which sub-pixel accurate detections are obtained (Fig. 1b,c). Both tasks can be naturally extended to an arbitrary number of dimensions, allowing Spotiflow to model 3D volumes directly and efficiently without any need of extensive postprocessing. Our method generalizes well to unseen samples and removes the requirement of manual threshold tuning in typical end-to-end iST workflows. Spotiflow is available as an easy-to-use Python library and as a napari [21] plugin (Supp. Video 2).

## Results

### Deep stereographic flow regression

To compute spot coordinates from a given microscopy image, Spotiflow uses a convolutional neural network (U-Net [22]) that is trained to predict two distinct but synergetic targets: *Gaussian heatmaps* and the *stereographic flow* (Fig. 1b, Supp. Fig. 1). For simplicity, we focus on the 2D case, but we note that the extension to *n* dimensions is straightforward (see Methods for the 3D case). The first target, *Gaussian heatmaps[23]*, are real-valued images of different resolutions in which each pixel can be interpreted as the probability of that position being a spot center (Fig. 1c, Supp. Fig. 1, Supp. Note 3.1). We predict a multi-scale hierarchy of heatmaps by processing their respective network decoder feature maps, which jointly contributes to the optimized training loss. We found this approach to be beneficial for training convergence, especially when only few spots are present (Methods, Supp. Fig. 1, Supp. Note 1). The second target, which we denote *stereographic flow*, is an alternative representation of the closest-spot vector field that, for every position, points to the closest spot. The stereographic flow is defined as the inverse stereographic projection of the n-dimensional local offset vector field in ℝ^*n*^ onto the *n*-sphere *S*^*n*^ ⊂ ℝ^*n*+1^. Crucially, this embedding maps all offsets for points far away from spot locations to a common value (specifically, the south pole of the unit sphere in the 2D case) therefore avoiding the problem of indeterminate offset prediction for distant locations (Fig. 1c, Methods, Supp. Fig. 2, Supp. Note 3.1, Supp. Video 3). To produce the final spot coordinates from a given prediction, we use the peaks of the highest-resolution heatmap to obtain preliminary spot locations, which we refine with the inverted stereographic flow (Methods), allowing modelling of subpixel locations and substantially lowering localization errors (Supp. Table 1, Supp. Fig. 3).

**Figure 2:**
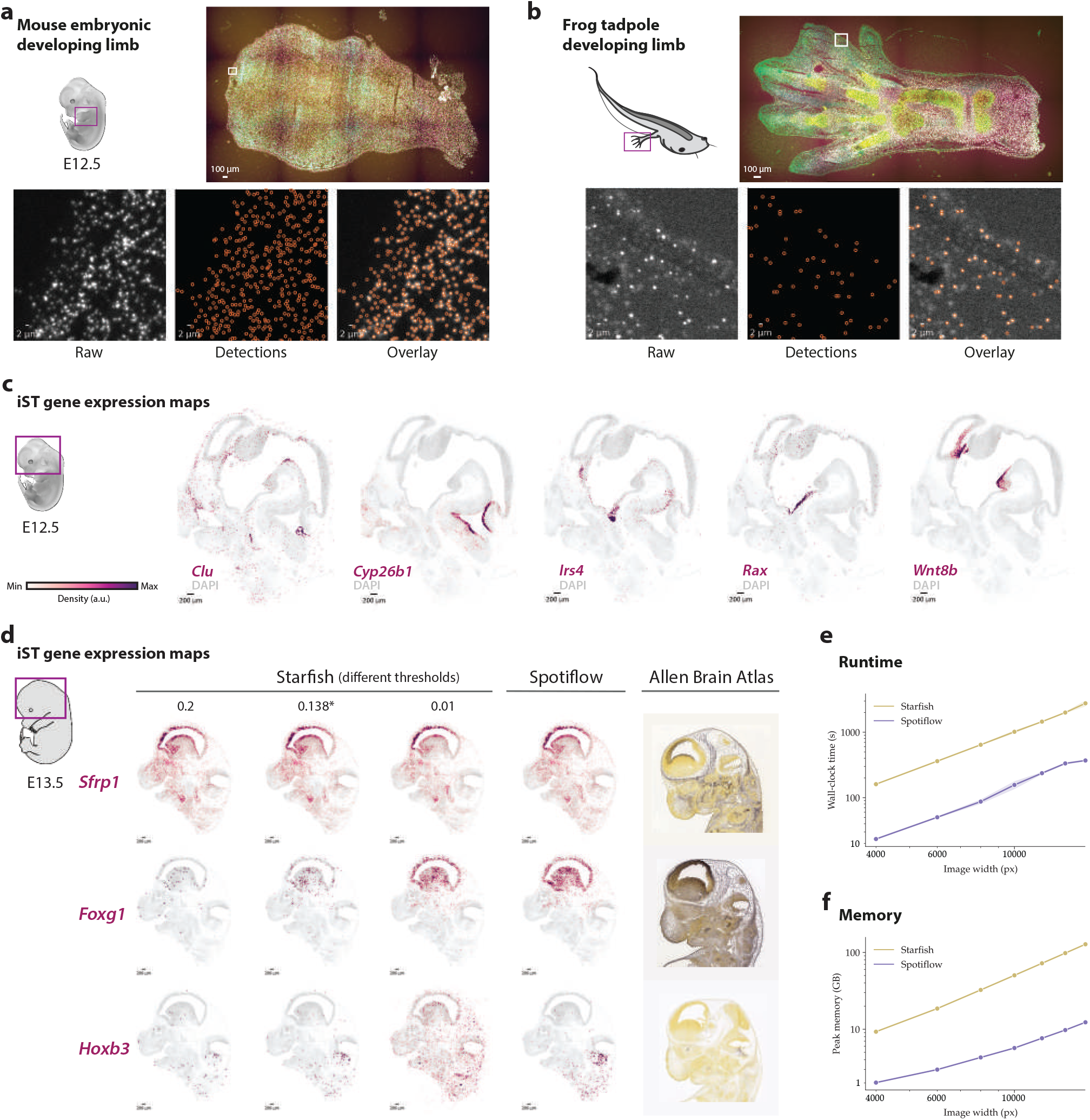
Application of Spotiflow in a variety of imaging-based transcriptomics settings. **a, b)** Predictions of a pretrained Spotiflow model on two out-of-distribution samples, a mouse embryonic developing limb (a) and a frog tadpole developing limb (b). The Spotiflow model was trained on the *HybISS* dataset, consisting only of images of embryonic mouse brain embryos. **c)** Gene expression maps based on Spotiflow of an El 2.5 mouse embryo brain processed using HybISS. Five different genes (*Clu, Cyp26b1,lrs4,Rax, Wnt8b*) involved in neurodevelopment overlaid on theDAPI channel are displayed. The Spotiflow model used was trained on the *HybISS* dataset. **d)** Comparison of gene expression maps based on Spotiflow *vs*. the default LoG detector in Starfish [14] of an El 3.5 mouse embryo brain processed using HybISS. Depicted are results for three different genes (Sfrp, *Foxgl*, HoxbT)-The Starfish detector is run at three different thresholds (0.2, 0.01 and 0.138, the latter being the optimal on the *HybISS* training dataset) as well as the Spotiflow model trained on *HybISS* (using the default threshold). The last column contains an ISH reference of similar sections from the Allen Brain Atlas (ISH) for the three depicted genes. **e, f)** Runtime (e) and memory (f) assessment of both methods in an end-to-end setting. Depicted are wall-clock time (e) and peak CPU memory usage (f). Memory usage excludes constants such as library imports or interpreter initialization.

**Figure 3:**
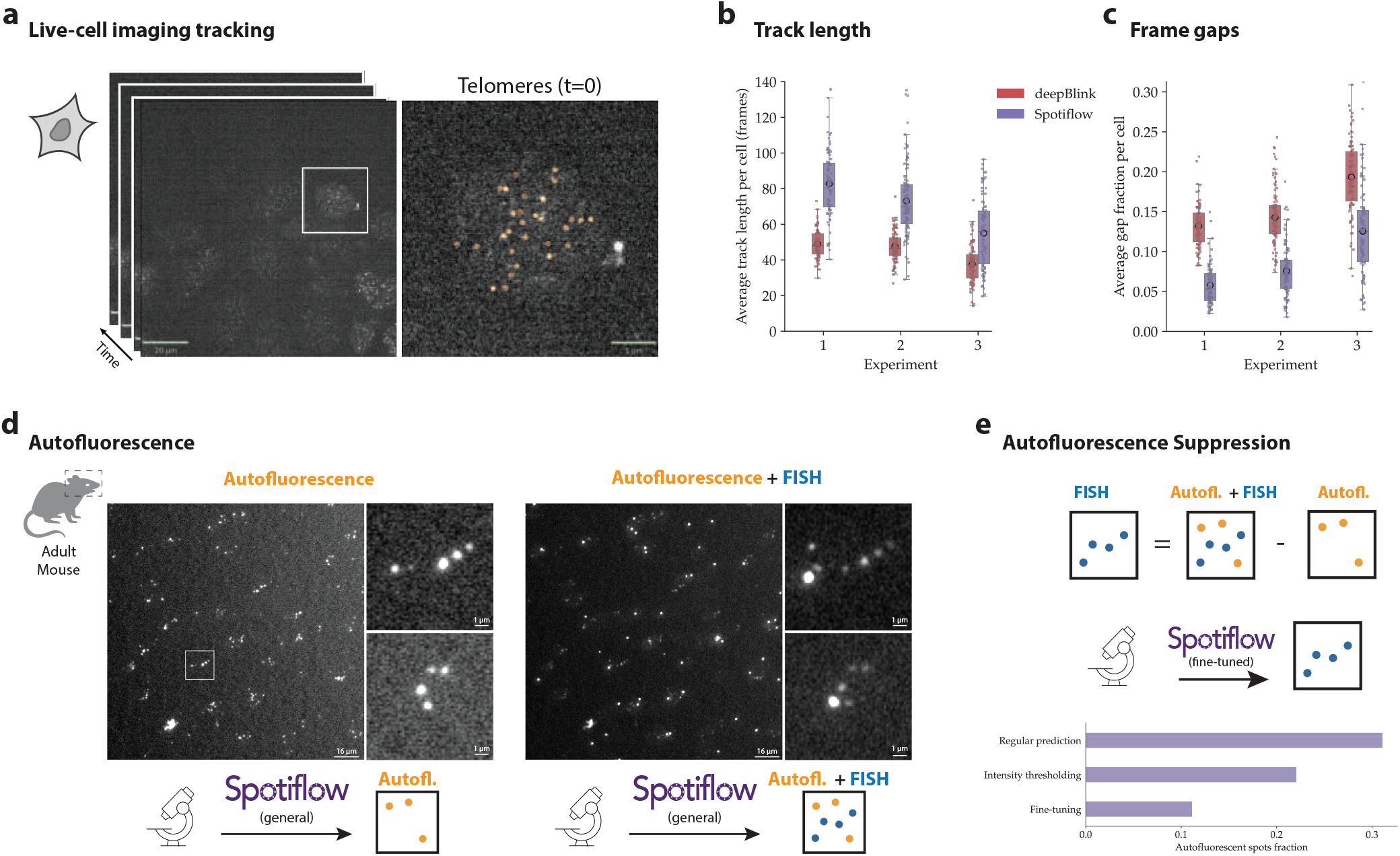
General applications of Spotiflow: live-cell imaging and autofluorescence-aware training, **a)** Live-cell acquisition of HeLa cells with labeled Telomeres (orange). **b)** Quantification of Telomere track length on three different experiments using deepBlink and Spotiflow to detect spots per frame, which are then tracked using TrackMate [25]. Telomeres are expected to be stable throughout the movie, thus longer tracks are expected. **c)** Quantification of number of frames where a track does not contain any detected spot (gap fractions). Smaller gap fractions indicate more stable detections. **d)** Depiction of setting where an autofluorescent cycle of an adult mouse brain is obtained previously to an hybridization-based iST protocol of the same tissue. **e)** Top: non-autofluorescent spots can be isolated by subtracting the spots detected in the autofluorescence round from all detections in the FISH signal. Middle: the same Spotiflow model used to retrieve the detections is then fine-tuned to detect only non-autoflorescent spots. Bottom: barplot depicting the amount of autofluorescence detected (lower is better) on real test data before (*baseline*) any removal, after intensity-based removal (*Intensity thresholding*) and after fine-tuning (*fine-tuning*). For a fair comparison, the intensity threshold for *Intensity thresholding* was chosen so that the amount of non-autofluorescent spots is exactly the same as for *fine-tuning*.

### Spotiflow achieves superior accuracy across imaging domains

We systematically assessed the performance of Spotiflow on multiple datasets in comparison with other commonly used methods. Specifically, we compared against the multi-scale Laplacian-of-Gaussian blob detection (LoG/starfish) implementation [24] used in the popular *iST* framework *starfish* [14], Big-FISH [15], the radial symmetry-based method RS-FISH [16], and the deep learning-based method deep Blink [19]. We first generated two synthetic datasets of diffraction-limited spots (Fig. 1d): one using a simple Gaussian PSF model (*synthetic-simple*), and another using a more realistic image formation model including autofluorescence and optical aberrations (*synthetic-complex*, Methods). We found that on *synthetic-simple* all methods achieved close to perfect scores (*F*_1_-score of 0.967–0.995), which is expected due to the limited complexity of the simulated images (Fig. 1d, Supp. Table 4). However, on the more realistic *synthetic-complex* dataset, classical methods showed a substantial performance drop (*F*_1_ = 0.758 – 0.836) while Spotiflow achieved the best detection performance (*F*_1_ = 0.929) followed by the other deep learning-based method (deepBlink, *F*_1_ = 0.915). This demonstrates the advantages of learning approaches for more complex datasets and highlights the limitations of overly simplistic simulations in benchmark scenarios. Similarly, when generating images at different noise conditions, spot densities and mimicking dense clusters, we found Spotiflow to consistently outperform other methods, demonstrating its effectiveness across a wide range of adverse imaging conditions (Supp. Fig. 4 & 5, Supp. Table 6).

**Figure 4:**
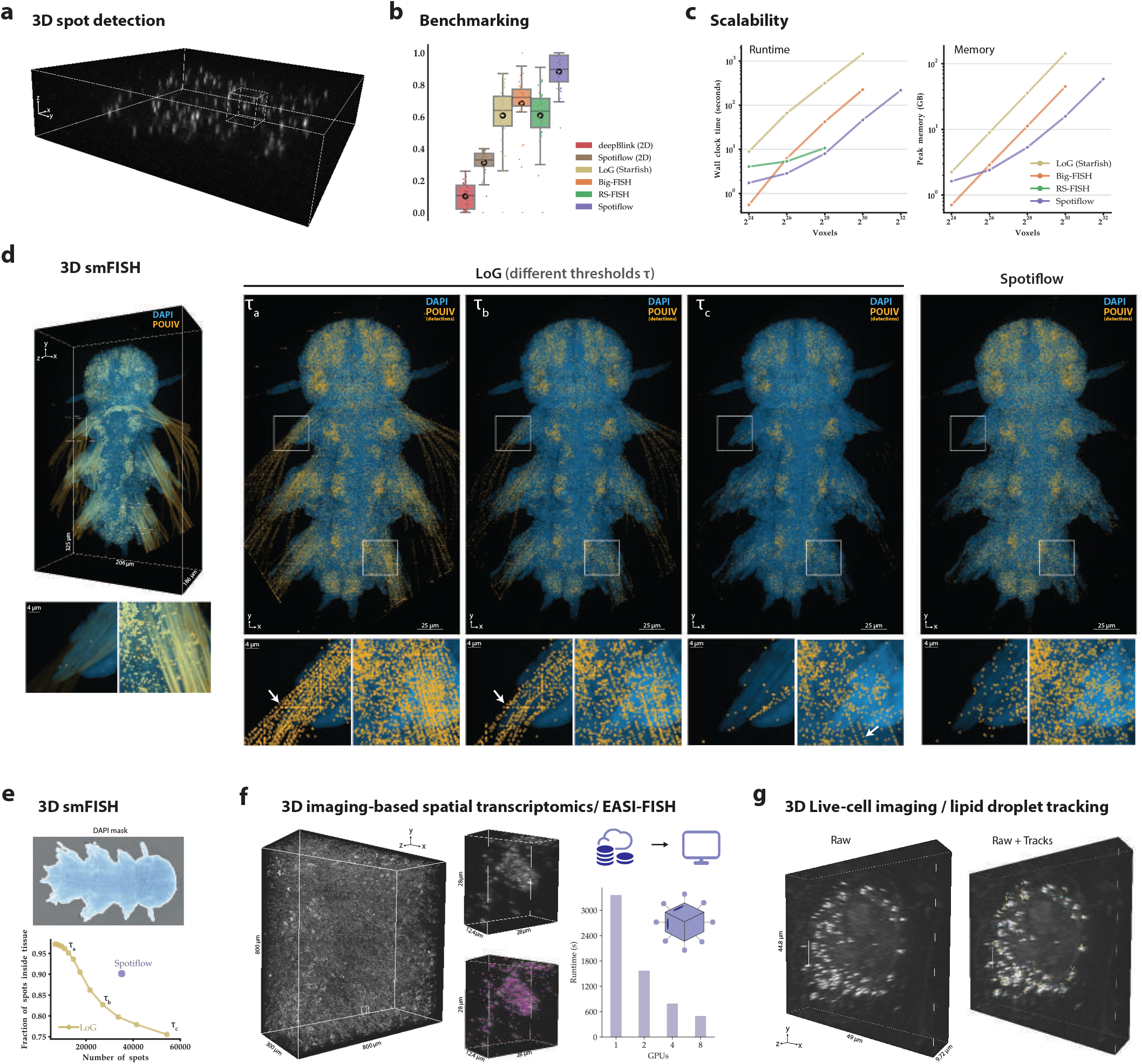
Assessment of spot detection performance on volumetric data with Spotiflow. **a)** Sample stack of the *synthetic-3D* dataset, which we use for benchmarking. **b)** Benchmarking results for different methods on the dataset *synthetic-3D*. Learning methods with the suffix (*2D*) were run per plane (YX) independently and then merged together to obtain 3D detections with the procedure implemented by [19]. deepBlink 3D is not shown as it does not offer a native 3D implementation. F_1_ [3] is depicted. **c)** Time (left) and memory (right) scalability assessment of native 3D methods. Depicted are wall-clock time (in seconds, left) and peak CPU memory usage (in gigabytes (GB), right). Memory usage excludes constants such as library imports or interpreter initialization. **d)** Spot detection comparison on a 3D smFISH volume of *Platynereis dumerilii*. A small amount of subvolumes were annotated, which were then used to optimize the parameters of LoG or fine-tune a Spotiflow 3D model. In the 3D view (left), DAPI is shown in blue and raw *Brn3* signal in orange. The Z-projections depict LoG detections at different thresholds *τ* as well as Spotiflow detections (orange). **e)** Quantification of ratio of number of detections inside the tissue. Semantic segmentation was performed on DAPI in order to obtain a tissue mask. **f)** Results of applying Spotiflow 3D to a large EASI-FISH stack [13]. The 3D views display an overview and an inset (without and with Spotiflow detections). A depiction of Spotiflow distributed capabilities is shown on the right, including runtime analysis using different numbers of GPUs. **g)** Lipid droplet tracking with Spotiflow and TrackPy on a live-cell label-free volumetric movie. The Spotiflow model used was pre-trained on *synthetic-3D* and fine-tuned on two frames of another movie.

We next assessed Spotiflow on several publicly available and in-house generated manual annotated datasets from multiple iST modalities (DNA FISH, MERFISH, HybISS, smFISH, *cf*. Supp. Fig. 6, 7, 8, Supp. Table 2, 3). We observed that Spotiflow again outperforms all other methods, including deepBlink, achieving a consistently high detection rate and localization accuracy (assessed with the Fl score and the *Panoptic localization quality PLQ* respectively, *cf*. Methods) on all modalities (Fig. 1d, Supp. Fig. 9, Supp. Note 3.3). The performance difference to threshold-based classical methods is particularly prominent for the HybISS and MERFISH datasets, which contain substantial background signal (*F*_1_ = 0.796/0.861 *vs*. 0.531/0.790 for *e.g*. LoG-starfish). Interestingly, a Spotiflow model jointly trained on all diverse datasets (*general*) achieves an almost equal performance to models trained on individual datasets, demonstrating the inherent capacity of Spotiflow to capture diverse image characteristics in a single model. Spotiflow also exhibits a higher level of robustness to common artifacts found in microscopy images such as patterned noise, hot pixels, and vignetting effects when compared to LoG (Supp. Fig. 10), as well as to border effects due to common image transformations (Supp. Fig. 11).

We evaluated the utility of Spotiflow for non-iST modalities by annotating two datasets from single frames of live-cell recordings of HeLa cells with labeled telomeres and the telomeric repeat containing RNA *TERRA* (Methods). As before, Spotiflow outperforms all other methods both in detection quality *F*_1_ and localization quality PLQ (Fig. 1d), demonstrating the general applicability of our method beyond the iST domain. In relation to the training data requirements of the model, we observed that fine-tuning based on synthetic data can substantially reduce the annotation requirement for novel out-domain datasets, quickly approximating the accuracy of our benchmark training dataset composed of hundreds of annotated 512 × 512 images. Specifically, when we fine-tuned a Spotiflow model initially pre-trained on *synthetic-complex* with an incrementally increasing number of out-domain training images from the live-cell dataset *Terra*, we found that already as few as four training images sufficed to achieve good accuracy (*F*_1_-score 0.738 *vs*. 0.174 when training from scratch, Fig. 1f, Supp. Fig. 12). Similarly, when fine-tuning the *general* model on only three 930 × 1306 *DNA FISH* images we achieved excellent performance on the full test set (*F*_1_-score 0.92 *vs*. 0.77 pre-fine-tuning) despite the little training data used. We also assessed the dependency of detection performance on the amount of annotation mistakes that might be present in the training data. To that end, we trained models with data that was artificially perturbed with varying amounts of annotation noise such as dropping annotations and introducing random annotation shifts (Supp. Fig. 13). While as expected the localization accuracy PLQ decreases steadily with the amount of introduced displacements (Δ*PLQ* ∈ [–0.15, 0], Supp. Fig. 13c-d), we found that the *F*_1_ score of Spotiflow models on an unperturbed test set remains largely unchanged even when dropping as much as half the training spots or introducing random shifts of 1px (Δ*F*_1_ ∈ [–0.02,0], Supp. Fig. 13a-b), highlighting the robustness of the detection performance to annotation noise. These results underscore the efficiency of Spotiflow models in adapting to different modalities and out-of-distribution (OOD) samples with minimal annotation, facilitating rapid adoption by end-users.

### Spotiflow produces robust and accurate gene expression maps in iST

We next investigated the generalizability of the 2D pre-trained models to the variability in sample types, which can encompass differing signal-to-noise ratios, unique artifacts, and distinct background features. We trained a Spotiflow model on HybISS-processed mouse embryonic brain sections (corresponding to the *HybISS* benchmarking dataset) and applied it on a variety of out-of-distribution HybISS samples originating from different tissues and probesets (mouse embryonic limb, frog tadpole developing limb, mouse gastruloid, radial glia progenitor cell cultures). Even though these images exhibit noticeably different structures with varying backgrounds and contrast compared to the training images, we found that the pre-trained model yielded qualitatively excellent transcript detection results without the need of any threshold-tuning (Fig. 2a,b and Supp. Fig. 14).

We then assessed the impact of the increased robustness and accuracy of Spotiflow for a full end-to-end iST experiment (Methods) by using a starfish gene decoding pipeline where we swapped the spot detection component from the default LoG detector to Spotiflow. We processed sections of developing mouse brains at different timepoints, E12.5 (Fig. 2c) and E13.5 (Fig. 2d), using HybISS to spatially resolve 199 genes involved in neurodevelopment (Methods, Supp. Table 5). The resulting gene expression maps obtained with Spotiflow show a gene-dependent spatial pattern that is consistent with previous results (Fig. 2c, Fig. 2d, Supp. Fig. 15). While for intensity-based methods (*e.g*. LoG/starfish) the quality of the obtained gene expression maps is highly sensitive to the used threshold and thus requires channel-specific threshold choices, Spotiflow is threshold agnostic and does not require any manual tuning (Fig. 2d, Supp. Fig. 15). In addition, we found that in this end-to-end *iST* setting, Spotiflow is an order of magnitude more time and memory efficient than the default starfish pipeline, especially for large images (Fig. 2e,f).

### Spotiflow extends beyond single image spot detection

To demonstrate Spotiflow’s impact on tasks beyond single image spot detection, we next applied it to a single-molecule detection and tracking task. Here, we considered a dataset (*TERRA*) where both telomeres and noncoding RNA molecules where recorded in live HeLa cells along a time lapse experiment (Methods, Fig. 3a). The resulting image set presents different challenges compared to iST images, such as photobleaching causing the temporal decrease of image contrast, non-specific dot-like structures inside the cell nucleus, and unspecific signal that can lead to erroneous and unrealistic short tracks (Fig. 3a). After detecting spots with Spotiflow, we tracked them using TrackMate [25] (Methods). We also detected spots with deepBlink and tracked them for comparison. For both telomeres and *TERRA*, the robustness of Spotiflow’s detections at changing imaging conditions led to longer, more consistent (gap-free) tracks compared to deepBlink (Fig. 3b,c, Supp. Fig. 16, Supp. Video 4), demonstrating the significant impact on the estimates of biological parameters our method is able to provide.

We next hypothesized that the content-awareness of Spotiflow could be leveraged to solve tasks which are infeasible for classical spot detection methods. To explore this we examined whether Spotiflow could effectively differentiate between transcript-derived spots and spot-like patterned autofluorescence structures, such as those from lipofuscin [26, 27], that often render data collected from adult brain unusable. Applying Spotiflow to a HybISS-processed adult mouse brain section with a specific bootstrapping scheme (Methods, Fig. 3d,e) we achieved a 2x decrease in the number of autofluorescent spots detected in one channel (Fig. 3e) compared to intensity-based spots removal. This is particularly notable considering that such a discrimination task is challenging even to experienced human annotators. Finally, we note that Spotiflow can be trained to localizing non-spot like structures and extended objects. As an example, we applied Spotiflow to a public dataset of microbial colonies cultured on agar plates ([28], Supp. Fig. 17) demonstrating the generality of our approach.

### Spotiflow yields robust and scalable gene localization in 3D

We next assessed the 3D version of Spotiflow, which uses 3D convolutions and volumetric extensions of the two target tasks described before to natively allow prediction on volumetric data. To quantitatively benchmark performance we created realistic synthetic volumetric data (Fig. 4a, Methods) and compared against both against 3D versions of the classical methods (LoG, Big-FISH, RS-FISH) as well as the 2D deep learning methods using the Z-blending strategy proposed by the authors of [19]. For Spotiflow, we included both the native 3D model (simply Spotiflow) as well as a 2D model using the same Z-blending strategy (spotiflow (*2D)*). Overall Spotiflow achieves the highest score (*F*_1_ = 0.882), with a substantial advantage compared to the second-best performing method (*Big-FISH, F*_1_ = 0.683, Fig. 4b, Supp. Table 7). Notably, the 2D learning-based models with a blending strategy clearly underperform (*F*_1_ = 0.102 and *F*_1_ = 0.311) compared to a native 3D method (Fig. 4b, Supp. Table 7), highlighting the benefits of native volumetric prediction. Additionally, we show that Spotiflow allows effective processing of blurry iST data that would otherwise require deconvolution. Using the synthetic 3D iST dataset from [29], we found that applying the pretrained Spotiflow model directly to raw images yields a higher fraction of correctly identified spots than deconvolution followed by an intensity-based method, irrespective of the spot density (*e.g*. 97% *vs*. 91% for 1600 spots per stack, Supp. Fig. 18). Notably, as in the 2D case Spotiflow processing time scales favourably with volume size *e.g*. taking less than 4 min for a stack of size ∼ 1600^3^ voxels for which other methods struggle on the same hardware (Fig. 4c).

To evaluate the performance on real volumetric data, we next used Spotiflow to detect *POUIV* transcripts in smFISH volumes of *Platynereis dumerilii* (Fig. 4d). *P. dumerilii* samples can exhibit highly intense and structured autofluorescence patterns at specific wavelengths, posing a challenge for intensity-based methods. We first pre-trained a Spotiflow model on the synthetic volumetric data and then fine-tuned the model on a few manually annotated subvolumes of size ∼ 256^3^ (9 for fine-tuning, 6 for validation) of a different sample. In order to appropriately account for the strong autofluorescence we included two crops of unlabeled autofluorescent regions to the fine-tuning data, highlighting the capability of the method to achieve performance gains with only minor annotation burden (Fig. 4d). Compared to the intensity-based method *LoG*, Spotiflow qualitatively produced less autofluorescent detections (Fig. 4d, Supp. Video 5) for a comparable amount of spots. To quantify this observation we computed the fraction of detections inside the tissue area *vs*. the total number of spots as a proxy of biological meaningful detection efficiency, finding that Spotiflow achieved a higher in-tissue efficiency for the same total number of detections compared to *LoG*, irrespective of the chosen *LoG* threshold (Fig. 4e, Methods).

We further applied Spotiflow to a 159 GB EASI-FISH volume of a lateral hypothalamus section of a mouse brain [13] in order to check transferability and assess the scalability of the method. In this case, the model used before for 3D smFISH data was able to accurately detect spots without further fine-tuning, highlighting the generalization capabilities of the method (Fig. 4f, Supp. Fig. 19). Moreover, running Spotiflow on the whole stack took little less than an hour (Fig. 4f) on a single GPU, a similar runtime to the fast Spark implementation of RS-FISH reported in [16]. By distributing the prediction across a commonly used multi-GPU compute node (8xA100, Fig. 4f, Methods), we achieved a runtime of less than 10 minutes thus demonstrating fast inference on large volumes in scenarios where GPU compute clusters are available.

We finally assessed whether Spotiflow can be used on a different domain by tracking lipid droplets in label-free microscopy volumetric movies (Fig. 4g, Supp. Video 6) of patterned COS7 cells. Similarly to the 3D smFISH case, we annotated the first two frames of one 3D movie in order to fine-tune a Spotiflow model that had been pre-trained on synthetic data. This model was then successfully applied to an out-of-training 3D movie, with Spotiflow being able to accurately detect lipid droplets despite the little fine-tuning data used, allowing successive tracking with TrackPy [30].

### Spotiflow is an efficient and user-friendly method for 2D and 3D spot detection

Training a Spotiflow model is fast even on modest resources (∼lh on a single NVIDIA RTX 4090 GPU for a 2D model) and our implementation based on PyTorch[31] is an order-of-magnitude faster and over three times more memory efficient than commonly used methods especially for larger images, with *e.g*. prediction time of 80s for an image of size 32*k* × 32*k vs*. 1000s for LoG/starfish (Fig. 1e, Supp. Note 3.4). To facilitate the adoption by end-users we provide an easy-to-use API (Fig. 5a, Supp. Fig. 20) and a command-line interface (Fig. 5b), extensive documentation, distribute Spotiflow as an easy-to-use napari[21] plugin (Fig. 5c), and provide several pre-trained models that can be used out-of-the-box for a variety of datasets (Supp. Video 2). We also offer our implementation for easily parallelizing across multiple GPUs during inference, which enables quasi-linear speed-up when using multiple GPUs (Fig. 4e), as well as allow predicting on Zarr data on cloud storage (*e.g*. AWS S3), thus allowing for fast and scalable processing on modern computing environments. Finally, we also provide an integration with the popular software *Starfish*[14] which eases the addition of Spotiflow to existing iST pipelines, both for 2D and 3D (Fig. 5d).

**Figure 5:**
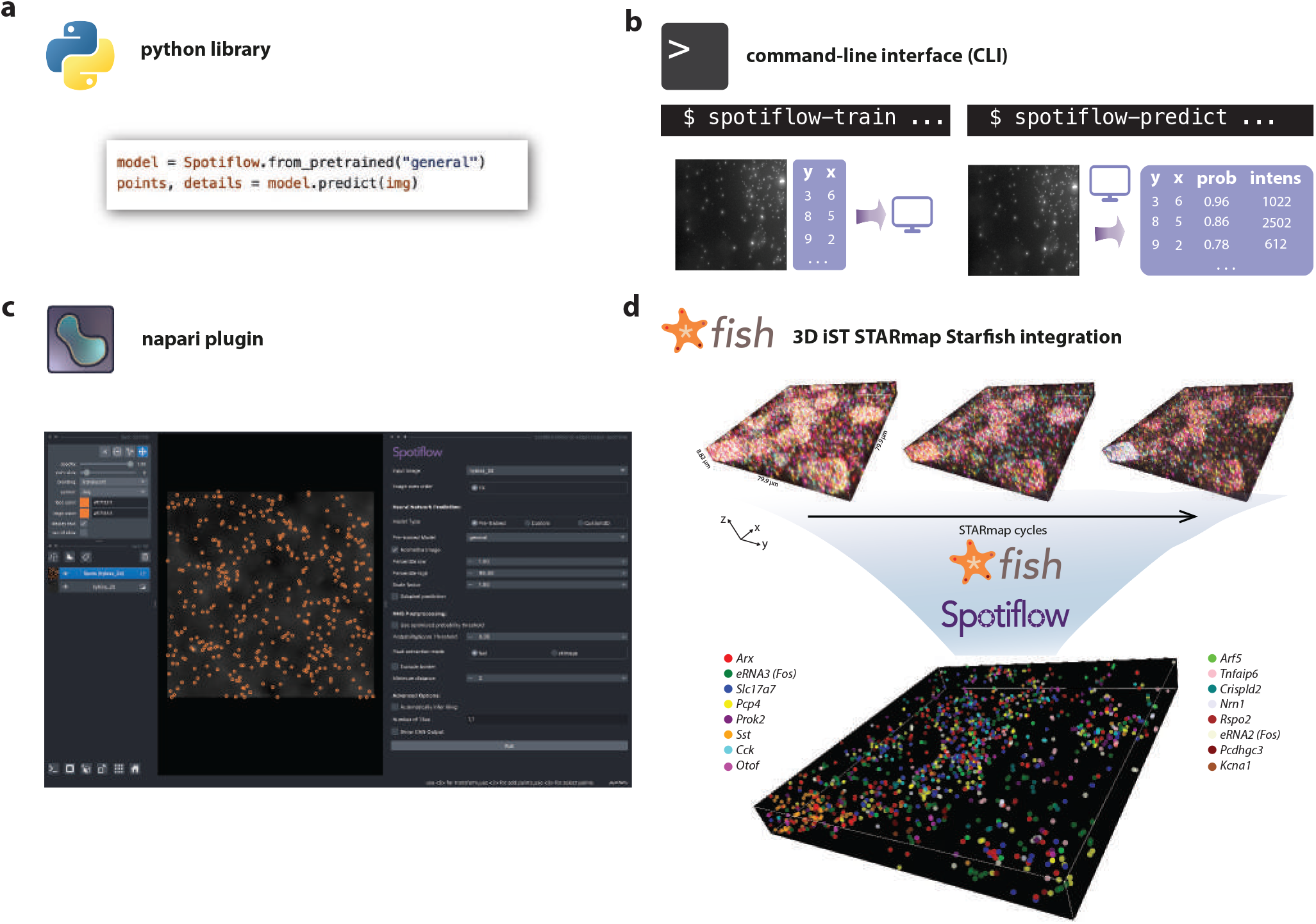
Spotiflow accessibility and integration. **a)** Spotiflow is usable through a simplified Python API, which also allows loading pre-trained models. **b)** Overview of the napari plugin of Spotiflow, offering pre-trained models for inference as well as a user interface for training and fine-tuning models. **c)** Demonstration of Spotiflow in an end-to-end 3D imaging-based transcriptomics setting. Top: field-of-view of three STARmap 4-channel cycles of a mouse brain volume [12]. Middle: Spotiflow can be plugged effortlessly into an existing Starfish [14] pipeline to enhance the spot detection task. Bottom: transcripts corresponding to 16 different genes obtained with Starfish and Spotiflow.

## Discussion

We presented Spotiflow, a threshold-independent and scalable 2D and 3D spot detection method that provides high-quality detections across a variety of real iST and non-iST modalities. Spotiflow’s ability to generalize to out-of-distribution samples reduces the need for re-training in new acquisition settings and its threshold-agnostic nature simplifies end-to-end iST workflows by eliminating the need for channel and round-specific manual threshold tuning that is currently often required to achieve optimal performance. These features support the construction of unified pipelines and standardized processing routines, which is crucial in the current state of development of the field, with different laboratories evolving protocols and acquiring iST data with different imaging setups. In cases where a pretrained model might be insufficient, fine-tuning on a small number of annotated images allows adapting Spotiflow to new imaging conditions, reducing the annotation burden for end-users. Spotiflow’s computational efficiency, being substantially faster and more memory-efficient than other techniques, enables streamlined processing of gigapixel-sized images, as seen in many iST applications, and makes it ready to support extensive data collection campaigns to build organism-sized atlases. Our live-cell imaging experiments indicate Spotiflow’s flexibility to various microscopy modalities, and we foresee its broad utility to other imaging-based methods where localized structures need to be detected. Furthermore, Spotiflow achieves state-of-the-art performance in 3D, overcoming the bottleneck that is inherent to plane-wise application of 2D methods.

Despite these strengths, model performance depends on the quality of training annotations, which can introduce biases or may limit the ability to perform accurate subpixel localization. A potential approach to unlock very precise subpixel modelling with Spotiflow is to train a model on the results of fitting procedures, or use said fitting procedure to define initial coarse annotations. Additionally, fine-tuning may require extra annotations when imaging conditions or object characteristics differ significantly from the pretraining datasets, though our results demonstrate that the required annotation effort can be minimal.

Finally, we anticipate that the presented *stereographic flow* task will impact other areas of image analysis where the prediction of dense vector fields has been successfully applied (*e.g*. cell segmentation [32, 33]). Overall, Spotiflow offers a robust solution for spot detection in imaging-based spatial transcriptomics and light microscopy opening up new possibilities for high throughput and high resolution spatial analyses.

## Author contributions

A.D.M. developed the idea, wrote the software, performed computational experiments, annotated data, performed and interpreted analyses, created figures and wrote the manuscript. A.H.C. acquired HybISS data (mouse embryo, frog tadpole and gastruloids), helped interpreting the related analyses, annotated 2D data and created figures. I.K. designed the mouse brain gene panel, acquired HybISS data (mouse embryo), helped interpreting the related analyses, annotated 2D data and ran preliminary computational experiments. A.S. acquired HybISS data (adult mouse), helped interpreting the related analyses and annotated 2D data. E.K. acquired the 2D live-cell imaging movies, annotated 2D data, performed tracking and helped interpreting the related analyses under the supervision of J.L.. G.T. prepared limb samples and E.S. generated gastruloids under the supervision of C.A.. L.S. acquired the 3D smFISH sample of *P. dumerilii* under the supervision of D.A.. E.B. processed the raw 3D smFISH stack of *P. dumerilii*, annotated 3D data and helped interpreting the related analyses under the supervision of A.K.. B.D. acquired the live 3D movies under the supervision of S.M.. G.L.M. supervised the project, developed the idea, designed the mouse brain gene panel, interpreted analyses, created figures and wrote the manuscript. M.W. supervised the project, developed the idea, wrote the software, interpreted analyses, created figures and wrote the manuscript. All coauthors read and approved the manuscript.

## Data availability

The benchmark datasets, including images and annotated spots, are available at https://zenodo.org/records/14514463 [34].

## Code availability

Spotiflow is available as an open-source Python library and as a napari plugin at https://github.com/weigertlab/spotiflow.

## Acknowledgements

We thank members of the Weigert and La Manno labs as well as Lars Borm (KU Leuven) for their feedback and discussions of the project. We further thank Stephan Preibisch (HHMI Janelia) for providing access to the EASI-FISH data. We would also like to thank the EPFL BioImaging & Optics Core Facility (BIOP) and the EPFL Histology Core Facility for their assistance in imaging and sample preparation. This project was supported by the EPFL Center for Imaging. M.W. was supported by the ELISIR program of the EPFL School of Life Sciences and by generous funding from CARIGEST SA. L.S. and D.A. were funded by the Marie Sklodowska-Curie ITN “EvoCELL” #766053 and the ERC Advanced grant NeuralCellTypeEvo #788921. Work in J.L.’s group was supported by the Swiss National Science Foundation (SNSF) [310030_214833] and the SNSF-funded National Centre of Competence in Research RNA and Disease Network [205601]. E.K. was a recipient of a postdoctoral fellowship from the Peter and Traudl Engelhorn Stiftung [532515].

## Methods

### Spotiflow

#### Architecture overview

Given an input image and the corresponding spot center annotations, a U-Net [22] is trained to predict two different sets of outputs which encode the location of spots in the image: first *multiscale probability heatmaps*, and second, the *stereographic flow (cf*. Fig. 1b, Supp. Fig. 1). During training, the overall loss function optimized is

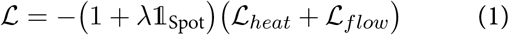

where ℒ_*heat*_ is the multiscale heatmap loss (see below), ℒ _*flow*_, the stereographic flow loss (see below) and 𝟙_spot_t is a pixel-wise indicator function which takes the value 1 if the pixel is very close to a spot location (whenever *Y* ^(0)^(*x*) ≥ 0.01 (see Equation 2 below) and 0 otherwise. *λ* ∈ ℝ is used to increase the loss contribution near spot centers (*cf*. Supp. Note 3.1). We set *λ* = 10, and *ε* ≈ 5px when training all our models.

#### Multiscale heatmap regression

Let *X* ∈ ℝ^*w*×*h*^ denote the input image and {*p*_*k*_} with *p*_*k*_ ∈ ℝ^2^ denote the ground truth spot center annotations. We first build a full resolution probability heatmap *Y* ∈ ℝ^*w*×*h*^ by generating a Gaussian distribution of variance *σ*^2^ centered at every spot so that the probability map decays around the annotated center in a Gaussian fashion.

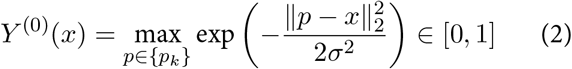

Note that instead of summing the individual Gaussian distributions, we take the maximum value at each pixel to create sharp boundaries between spots.

We further generate the heatmaps at *L* different resolution levels, where level *l =* 0 denotes the highest resolution and *I = L* — 1 the lowest. In order to generate a heatmap at resolution level *l* (*Y* ^(*l*)^) from *l* − 1 (*Y* ^(*l*—1)^),, we apply max pooling (with a downsampling factor of 2) to *Y* ^(*l*)^ and then process the result with a Gaussian filter of variance 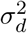 with a scaling prefactor of 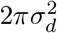, which effectively increases the variance of distributions and ensures the dynamic range of the heatmap is in the interval [0, 1] (*cf*. Supp. Fig. 1).

The U-Net backbone is then trained to regress all heatmaps 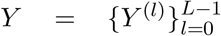 at the different scales (*multiscale heatmap regression, cf*. Fig. 1c, Supp. Fig. 1). We achieve this by adding a loss term at different stages in the decoder whose size correspond directly to the target to be regressed. More specifically, let *D*^(*i*)^, *i* ∈ [1, *L*],, denote the feature maps at the output of the *i*-th decoder stage in the U-Net. We process *D*^(*i*)^ with a lightweight convolutional module to compute the prediction 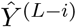 (*cf*. Supp. Fig. 1). A pixel-wise loss term is then computed between the ground truth heatmap *Y* ^(*l*)^ and the prediction 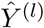 at every resolution level *I* with the binary cross-entropy loss. We then aggregate this terms in the overall objective function for the multiscale heatmap, ℒ

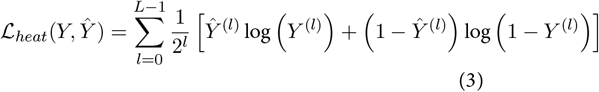

#### Stereographic flow

For each pixel (*i, j*) ∈ ℤ ^2^ of the image *X*, we first define a *local vector field V* = {*v*_*ij*_} = {(*v*_*x*_, *v*_*y*_) ∈ ℝ^2^} given by the vector from the pixel to the nearest ground truth spot (*cf*. Fig. 1b, Supp. Fig. 2). To induce stability and improve modelling at points far from spot locations, we make use of a scaled *inverse stereographic projection f* 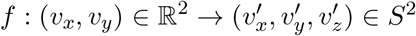 defined as

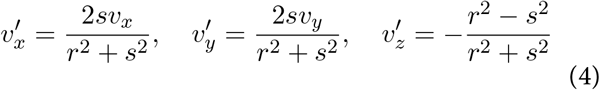

with 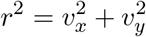 and 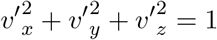
where *s* ∈ ℝ^+^ is a fixed length scale (we set *s* = 1). We define the *stereographic flow* 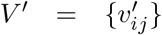 as the result of applying *f* to each component of the local vector field *v*_*ij*_. Effectively, we represent each element of the local vector field as a point on the unit 3D sphere *S*^2^ (note that this generalizes to arbitrary dimensions). In particular, *f* maps the zero vector (0, 0) to the north pole (0, 0, 1) and all vectors with infinite length (“points at infinity”) to the south pole (0, 0, −1). The stereographic flow is computed using an extra lightweight convolutional module operating at the highest resolution (*cf*. Supp. Fig. 1). The corresponding loss function is a pixel-wise weighted *L*_1_ loss ℒ_*flow*_ between the ground truth stereographic flow 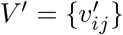 and the prediction 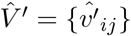:

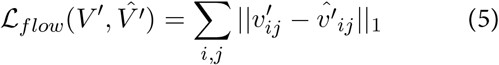

Note that, by construction, 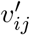 and 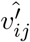 lie on the three-dimensional unit sphere *S*^2^ yielding a bounded target to be regressed.

Let *S*^’2^ = *S*^2^ \ {(0, 0, −1)} denote the set of all points in the unit sphere *S*^2^ but the south pole. The stereographic flow can be analytically inverted position-wise by applying the *stereographic projection* 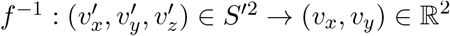:

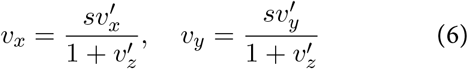

We note that despite the stereographic projection being undefined at the south pole (0, 0, −1), in practice we only invert the stereographic flow at positions that are close to a spot, which are embedded far from the south pole.

#### Inference

To retrieve the spot centers from the two outputs of the network (*i.e*. multiscale heatmaps and stereographic flow), we first detect all local maxima in the highest resolution predicted heatmap 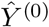. These local maxima are filtered so than only those above a specific threshold (*probability threshold t* ∈ [0, 1]) are kept. This threshold is optimized right after training on the validation data and thus does not need to be set explicitly during inference. This procedure results in a set of points {(*x*_*k*_, *y*_*k*_)} where *x*_*k*_, *y*_*k*_ ∈ ℤ.

These points are then refined using the stereographic flow to achieve subpixel precision by adding the corresponding predicted vector at every position. Specifically, let 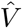 denote the pixel-wise stereographic projection of the predicted stereographic flow 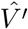, so that the vector at position 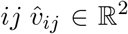. We generate the final set of points {*p*_*k*_}, which correspond to the spot centers, as 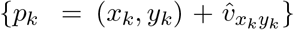, where {(*x*_*k*_, *y*_*k*_)} are the local maxima extracted from the full resolution heatmap and *p*_*k*_ ∈ ℝ^2^, thus allowing the prediction of non-integer (subpixel-precise) spot centers.

##### 3D architecture

The architecture of the 3D model follows the same principles as the 2D model (see above) but using 3D convolutions instead. The target tasks (*multiscale heatmap regression* and *stereographic flow*) are naturally extensible to an arbitrary number of dimensions by adapting accordingly. First, the multiscale heatmap is generated in 3D using the same procedure as described above using Eq. (2). The stereographic flow 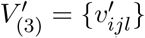 is obtained by applying the scaled inverse stereographic projection 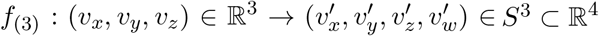, which is defined in a similar fashion to Eq. (4):

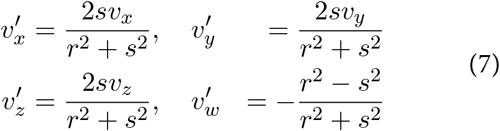

with 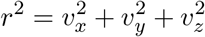 and 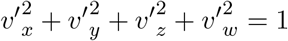.

As in the 2D case, *f*_(3)_ can be analytically inverted with the stereographic projection 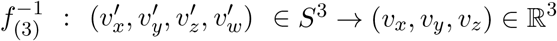

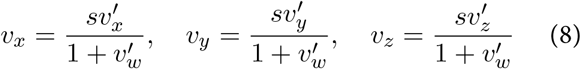

so subpixel-accurate locations can be computed.

While the model architecture is conceptually similar to the 2D counterpart, 3D computations are much more computationally expensive. To alleviate the workload and allow fast inference times on regular hardware, the 3D model allows predicting a downsampled version of the output (from now on, *gridding*) similarly to the 3D implementation of Stardist [35].

Let *h, w* and *d* denote the height, width and depth respectively of the input volume. Let *g* be the gridding (output downsampling) factor, which for simplicity we assume to be equal for every dimension. The input is first downsampled to size 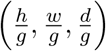 with a stem (specifically, a *g*-strided 3D convolutional module). The output of the stem is processed with the same backbone architecture as described before (a 3D U-Net), which yields a 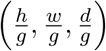 -shaped volumetric heatmap as well as a 4-dimensional stereographic flow of size 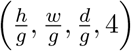 Note that the initial input downsampling operation carried through the stem is learnable, hence the neural network effectively operates on all pixels on the image.

The training must be “gridding-aware”, as it is inherent to the model architecture. In order to train the network, simply subsampling the targets would be suboptimal as it does not yield proper encoding of spots located at coordinates which are not multiples of *g* (from now on, *non-g-aligned*). Therefore, downscaled versions of the heatmap (*gridded heatmaps*) are produced with a slight variant of Eq. (2) which ensures that the pixel coordinates closest to non-*g*-aligned spots achieve the maximum possible value, which is 1. Moreover, the gridded stereographic flow is constructed so that non-*g*-aligned spots can be perfectly recovered, which can be thought as a built-in natural interpolation procedure. This ensures that coordinates other than multiples of *g* can be accurately retrieved without any loss of information as long as the distance between two spots is ≥ *g*. To illustrate with an example, let us assume that *g* = 2 and that there is a spot *p* at the location (3, 2, 15) in the original volume. Ideally, the heatmap peak localization (in the downsampled output space) is 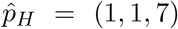 and the inverted stereographic flow at that position is 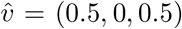. Thus, we can recover the original spot location as 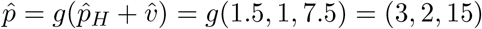. We note that gridding effectively brings down the ability of SPotiflow to separate spots that are very close together (closer than *g*). Nevertheless, we have found using *g* = 2 it to be a good tradeoff between performance and speed, obtaining considerable speedup (∼ 10x) at a marginal performance cost in most real settings. All results of Spotiflow 3D presented in the paper have been obtained with *g* = 2.

##### Spot detection benchmarking

#### Datasets (synthetic)

The dataset *synthetic-simple* was generated by randomly generating spot locations and placing Gaussian distributions with *σ* = 1.5 and varying intensity on a blank image, after which Poisson and Gaussian noise is added. The dataset *synthetic-complex* was generated similarly but instead of Gaussian spots we simulated realistically aberrated point-spread functions (PSFs) using the approach described in [36] and added fluorescence DAPI background, Gaussian noise, Perlin noise and Poisson noise at different levels yielding images with different SNRs across the dataset. Different densities (*i.e*. number of spots) were used to mimic different sparsity of real data. The dataset *synthetic-3D* was isotropically generated using the same procedure of *synthetic-complex*, but without the addition of DAPI fluorescence background.

#### Datasets (real)

We gathered the datasets *HybISS, Terra* and *Telomeres* by randomly cropping square tiles of width 512 and/or 1024 from different acquisitions described below (*cf*. sections *Data generation - imaging-based spatial transcriptomics* and *Data generation - Live-cell imaging (2D)*). The datasets *DNA FISH* and *MERFISH* were compiled by using raw images from [37] and [38] respectively, which we hand-annotated on *napari* [21]. In order to speed up the annotation process, we mostly resorted to use initial solutions obtained from LoG that we iteratively refined by adding, removing and/or moving the proposed spot centers using a custom annotation script. This script allows using Gaussian fitting to refine the integer positions obtained by LoG. For a reduced fraction of images (a subset of the datasets *HybISS* and *MERFISH*), we used an early prototype version of Spotiflow trained on different data instead of LoG to detect the initial candidates, which were then refined as described above. Different contrasts were considered when annotating to take into account potential uneven illumination. The annotated *smFISH* dataset was used as released in [19], which consists of a manually annotated subset of MERFISH data from [39].

#### Dataset preprocessing

Images were preprocessed equally for each method by normalizing them using percentile-based normalization:

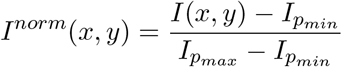

where *I*_*p*_ denotes the *p*-th percentile of the image intensity. We set *p*_*min*_ ∈ {1, 3} and *p*_*max*_ = 99.8 throughout our experiments. The same normalization procedure was performed in the 3D case. Note that the intensity values in the normalized images *I*^*norm*^ are not clipped to [0, 1].

#### Parameter tuning

Parameters specific to LoG (intensity threshold) and Big-FISH (variance of the filters) were optimized on the training split of each dataset. We did not optimize the intensity threshold on Big-FISH as the software has a custom threshold optimization procedure which works on an image-by-image basis. For RS-FISH, we optimized its parameters on the test split (thus overestimating its performance) due to the high computational load required and the large number of parameters that can be tuned (*cf*. Supp. Note 3.3). Learning methods (deepBlink and Spotiflow) were trained on the training split using their default configuration without performing any hyperparameter tuning (*cf*. Supp. Note 3.3). All reported scores are on the test split of the datasets.

#### Spotiflow (general) model

In order to assess the potential capacity of Spotiflow models, we trained the *general* model on a dataset gathered by merging all real 2D datasets (*HybISS, MERFISH, smFISH, Telomeres, Terra*) as well as the dataset *synthetic-complex*.

#### Metrics

To compute overall detection metrics for each image, we first uniquely match ground truth {*p*_*i*_} and predicted spots 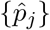 according to their spatial proximity via Hungarian matching [40]. We then define a spatial cutoff *c* ∈ ℝ and count a matched pair 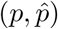 as *true positive* (TP) if their Euclidean distance 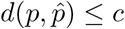, a predicted spot 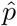 as *false positive* (FP) if there was no matched ground truth spot, and a ground truth spot *p* as *false negative* (FN) if there was no matched predicted spot. We then define the following metrics for each image

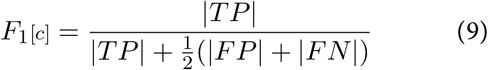

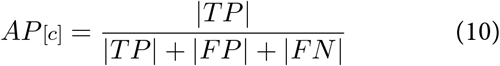

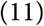

We also report the *F*_1_*AuC* [19] (*cf*. Supp. Note 3.2), which takes into account different spatial cutoffs:

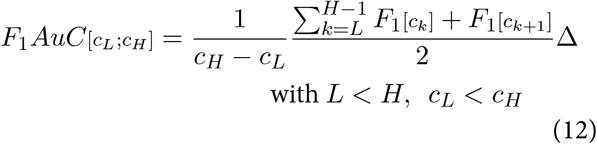

where Δ is a constant defined as *c*_*k*+1_ − *c*_*k*_ for any *k* ∈ [*L, H*). Finally, we adapt the Panoptic Quality segmentation metric [41], which incorporates the spatial localization accuracy, to the spot detection task. We refer to it as the *Panoptic Localization Quality* (PLQ):

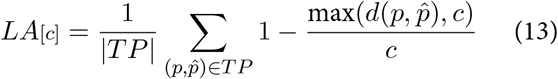

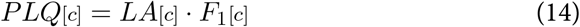

where *LA* is the localization accuracy (*cf*. Supp. Note 3.2). We report results at *c* = 3, *c*_*L*_ = 1 and *c*_*H*_ = 5.

#### Scalability assessment

In order to assess the scalability of different 2D methods (*cf*. Fig. 1e, Supp. Note 3.4), we used images of different sizes which were all obtained by consecutively expanding a center crop from a full HybISS cycle (*cf. Data generation - imaging-based spatial transcriptomics*). In the 3D case, a simulated stack from the dataset *synthetic-3D* (of original size 32×128×128) was replicated along each dimension to obtain larger volumes. The method deepBlink was not run on 3D as it cannot handle volumetric stacks natively.

Given the dependency of intensity-based methods on the number of spots, their parameters were set so that the number of detections were in the same order of magnitude across all methods. Elapsed time and memory consumption were obtained the Unix’s time module. Intensity-based methods were run on an AMD Ryzen Threadripper PRO 5965WX 24-Cores CPU with 256GB of memory. Learning methods (deepBlink and Spotiflow) were run on an NVIDIA GeForce RTX 4090 GPU (24GB).

#### Data generation

##### Imaging-based spatial transcriptomics

###### Tissue collection and preparation

All animal procedures were in accordance with the Swiss Federal Veterinary Office guidelines and as authorized by the Cantonal Veterinary Authorities and the Cantonal Commission for Animal Experimentation under the following licenses: cantonal animal license number VD3651 and national animal license number 33167 for mouse samples as well as cantonal animal license number VD3652c and national animal license number 33237 for frog tadpole samples.

###### Mouse embryos and frog tadpole samples

Mouse embryos at E12.5 and E13.5 were collected from wild-type cD1 pregnant mice dissecting out from the uterine horn in ice-cold PBS. Nieuwkoop and Faber (FB) stage 58 frog tadpole samples were collected in PBS. Immediately after collection, fresh tissues were cryopreserved in optimal cutting temperature (OcT) and stored at -80 ºc until sectioning. Tissues were sectioned with a cryostat (Leica cM3050 S) at 10 *µ*m, placed on SuperFrost Plus microscope slides, and stored at -80 ºc until HybISS processing.

###### Mouse gastruloid generation

Gastruloid generation was performed as previously described in [42]. Briefly, mouse embryonic stem cells (mEScs) (EmbryoMax 129/SVEV, gifted by Denis Duboule Lab) were cultured in gelatinized tissue culture dishes with 2i LIF DMEM medium consisting of DMEM + GlutaMAX (Gibco 61965-026) supplemented with 10% mES-certified FBS (Gibco 16141-079), non-essential amino acids (Gibco 11140-035), sodium pyruvate (Gibco 11360-039), beta-mercaptoethanol (Gibco 31350-010), penicillin/streptomycin (Gibco 15140-122), 100 ng ml-1 mouse LIF (EPFL Protein Facility), 3 *µ*M cHIR99021 (calbiochem: 361559) and 1 *µ*M PD0325901 (Selleckchem S1036). cells were passaged every 2-3 days and maintained in a humidified incubator (5% cO2, 37ºc). mEScs were collected after trypsin treatment, washed, and resuspended in prewarmed N2B27 medium (50% DMEM/F12 (Gibco 31331-028), 50% Neurobasal medium supplemented (Gibco 21103-049) with 0.5x N2 (Gibco 17502-048), 0. 5x B27 (Gibco 17504-044), non-essential amino acids (Gibco 11140-035), sodium pyruvate (Gibco 11360-039), beta-mercaptoethanol (Gibco 31350-010), 0.5x Glutamax (Gibco 35050-061) and penicillin/streptomycin (Gibco 15140-122). A total of 300 cells were seeded in 40 *µ*l of N2B27 medium in each well of a 96-well plate with a rounded bottom and low adherence (Thermo Fisher, 174925). Forty-eight hours after aggregation, 150 *µ*l of N2B27 medium supplemented with 3 *µ*M cHIR99021 was added to each well. A total of 150 *µ*l of medium was replaced every 24 h. Gastruloids were collected and flash-frozen 120 h after aggregation.

###### Radial glia progenitor culture

E11.5 mouse brains were collected from wild-type cD1 pregnant mice in ice-cold EBSS (14155-048, Life-technologies). Meninges were removed using fine-tipped forceps under a dissection stereomicroscope (Nikon SMZ18). Then, brains were fragmented into small pieces, transferred to a 50 ml plastic tube, and digested for 30-45 min at 37 ºc in 5 ml of solution containing 1mM cacl2, 1 mM Mgcl2, 100 U/ml of DNAse I (LS02058 Worthington, Lake Wood NJ), and 20 U/ml of previously activated papain (Sigma L2020). After that, the cell suspension was briefly decanted, transferred into a 15 ml plastic tube, and centrifuged at 300 rcf for 5 min at 4ºc. Then, the cells were resuspended in 3 ml of EBSS, and the suspension was transferred into a 15 ml plastic tube containing 3 ml of papain inactivating solution containing 50% FBS/50% EBSS. cells were centrifuged at 300 rcf for 5 min at 4ºc and then resuspended in culture media consisting of Neurobasal medium (21103049, Life Technologies) supplemented with L-glutamine (Gibco; cat. no. 25030-123), B27 (Gibco; cat. no. 17504-044), Gentamicin (15750037, Life Technologies), and 20 ng/ml of Epidermal growth factor (EFG, PeproTech catalog no. AF-100-15). Finally, cells were seeded on 8 well chamber slides (80841, IBIDI) and incubated in a humidified 37ºc incubator at 5% cO2 for 48 hours until HybISS processing.

###### In-situ sequencing by HybISS

To process all samples (apart from the adult mouse brain, *cf*. paragraph below) HybISS [9] was performed as published at protocols.io [43]. For embryonic mice, target genes were selected based on marker genes with expression within the target regions at the target developmental stages. Samples were imaged either on a Leica DMi8 epifluorescence microscope equipped with LED light source (Lumencor SPEcTRA X, nIR, 90-10172), scMOS camera (Leica DFc9000 GTc, 11547007) and 20x objective (Hc Pc APO, NA 0.8, air) yielding a pixel size of 0.34 *µ*m, or on a epifluorescence microscope Nikon Eclipse 90i equipped with LED light source (Lumencor SPEcTRA X, nIR, 90-10172), cMOS camera (Nikons DS Qi2) and 20x objective (cFI PLAN Pc, NA 0.75, air) yielding a pixel size of 0.15 *µ*m. In both microscopes, samples were imaged with 10% overlap between tiles to cover the entire tissue and between 8 and 12 z planes were acquired with 1 µm spacing among them. A full experiment results in a multicycle, multichannel image stack (5 cycles and DAPI + 4 HybISS signal channels) image stack.

###### In-situ sequencing by HybISS (adult mouse brain)

To process the adult mouse brain used for the autofluorescence removal experiment (*cf*. Fig. 3d,e), HybISS was performed on fresh frozen 6 weeks old mouse brains 10 *µ*m sections, using a Phi29 enzyme (NxGen F83900-1). Images were acquired using a 20X 0.8NA objective on a Zeiss AxioImager Z1 Wiedfield microscope with a PcO.edge 4.2bi camera. The microscope was controlled via MicroManager. Exposure times were, in order of acquisition: 3 ms for DAPI (Zeiss filter set 49: G365, FT 395, BP445/50), 450 ms for 750 nm (filters: Alluxa ultra 740.5-35 OD6, 766, 801.5-50 OD6), 300 ms for 650 nm (chroma BP 640/30, FT ZT640rdc, ET680/40), 300ms for 550 nm (filters BP546/12, LP T560lpxr, ET590-50), and 200 ms for 488 nm (chroma filters BP450-490, T510, ET 525/36). 130 tiles of 2048×2028 pixels with a 10% overlap were acquired to cover the entire tissue, with a pixel size of 0.3225 *µ*m. Each tile was a z-stack of 11 planes with a 0.8 *µ*m step size. Rounds of probe hybridization, imaging and stripping were performed with a modified Labsat microfluidics device from Lunaphore, allowing us to place the stainer chamber under the microscope. A quenching buffer (Lunaphore BU08) was used to reduce autofluorescence before bridge probe hybridization and an imaging buffer (Lunaphore BU09) was used during imaging.

#### Platynereis dumerilii 3D smFISH

##### Fixation of 6dpf P. dumerilii Larvae for in situ HcR RNA-FISH

*Platynereis dumerilii* worms were kept in continuous culture at EMBL Heidelberg. For details on P animal culture, see [44]. Every batch was generated by mating one pair of male and female adult worms inside a glass Becker with artificial sea water (ASW). Each batch was then incubated at 18ºc. At the 6th day post fertilization, batches were collected, poured together, and filtered with ASW in a cell strainer. Animals were incubated with 50 *µ*g/ml ProteinaseK in PTW (0.1%) Tween20 in PBS, DEPc treated) for 3’ at RT, washed in ASW, and anesthetized for 1’ in Noca2+-NoMg2+ ASW. Animals were then fixed in 4% PFA in PTW for 2h at room temperature (RT), followed by 5x 5’ washes in PTW. Animals were dehydrated in increasing concentrations of Ethanol (25-50-75% EtOH in PTW) for 5’ each, washed 4x 5’ with 1 ml Ethanol, and then stored for a maximumo of 6 months at -20ºc.

##### HcR RNA-FISH (v3.0) probe sets, amplifiers and buffers

HcRTM RNA-FISH (v3.0) DNA probes targeting the coding sequences of *P. dumerilii* POUIV and Prox genes were designed and synthesized by Molecular Instruments® (MI) (molecularinstruments.com). Probe sets included 14 and 20 probe pairs respectively. The reference transcript sequences can be found on GenBank at the following accession numbers: Pdu-POUIV, KC109636; Pdu-Prox, FN357281. The correspondent coding sequences used for the probe design are listed in the supplementary table. DNA HcR amplifiers, hybridization, wash and amplification buffers were purchased from MI. The probe sets, amplifiers and fluorophores combinations used for this experiment were the following: POUIV-B3_Amplifier-Alexa 546,Prox-B1_Amplifier-Alexa 647.

##### Whole-mount in situ HCR RNA-FISH (v3.0) of 6dpf P. dumerilii larvae

Animals stored in ethanol at -20°C were rehydrated at RT in decreasing alcohol concentrations (75–50–25% EtOH in PTW for 5’ each) and washed 3x 5’ in PTW. Samples were then permeabilized with 100 *mu*g/ml ProteinaseK in PTW for 3’, washed twice for 30’ in 20 mg/ml Glycine/PTW, and rinsed 3x in PTW. Animals were post-fixed for 20’ in 4%PFA in PTW, followed by 5x 5’ washes in PTW. The following is an adaptation of HCRTM RNA-FISH (v3.0) protocol of ML The original protocol with buffers composition can be found in [45]. Animals were transferred in 1.5 ml tubes and incubated in 200 *pi* HCR hybridization buffer (containing 10 mM VRC) at 37°C. After 30’ pre-hybridization, HCR probes, pre-diluted in 50 *µ*l HCR hyb. buffer, were added to the mixture at a final concentration of 4 nM in 250 *µ*l. Samples were hybridized over night (O/N) at 37°C (≅15–17h), shaking at 800rpm in a thermomixer. Samples were washed 5x 15’ in 1 ml HCR probe wash buffer at 37°C. Samples were then washed 3×5’ in 5XSSCT (0.1% Tween20 in 5XSSC) at RT. Samples were incubated in 100 *µ*l HCR amplification buffer at 25°C. In the meanwhile, HCR hairpins were heated in 0.2 ml PCR tubes at 95°C for 90” and then slow-cooled at RT in the dark for 30’. After 30’-lh of pre-amplification, HCR hairpins were diluted in 50 *µ*l HCR amp. buffer and added to the mixture at a final concentration of 60 nM in 150*µ*l. Samples were amplified O/N at 25°C (≅15–17h), shacking at 800 rpm in a thermomixer. Following amplification, samples were washed at 25°C in 1 ml 5XSSCT 2x 5’, 2x 30’, followed by a final 1 ml wash. Samples were either directly mounted for microscopy or stored in 5XSSCT at 4°C for a maximum of one week.

##### Autofluorescence quenching, nuclear counterstain, and mounting

HCR-stained samples were treated with 150 *µ*l Vector® TrueVIEW® Autofluorescence Quenching Kit for 2’ at RT, followed by 1 ml 5XSSCT washes, 2x 1’ and 2x 5’. Samples were then stained with 5 *µ*g/ml DAPI in 0,5 ml 5XSSCT for 30’, followed by 3x 5’ 1 ml 5XSSCT washes. Animals were mounted in ProLong™ Glass Antifade Mountant between two 1.5H coverslips glued together by 0.12 mm thick SecureSeal™ Imaging Spacer.

##### Microscopy and imaging parameters

Samples were imaged with a Leica TCS SP8 confocal microscope using an HC PL APO 40x/1.30 OIL CS2 objective. The zoom factor was adjusted to fit an entire larvae in the field of view with a pixel size of 129 nm and an image format of 2048 × 2048 px. Z-stacks of whole larvae were acquired with a z-step size of 0,5 µm. Three sequential by-frame scans using HyD detectors acquired fluorescence signals from DAPI, Alexa Fluor 546, and Alexa Fluor 647 dyes, excited by 405 nm, 561 nm, and 633 nm laser wavelengths, respectively. Mounting the larvae between two coverslips allowed the scanning from both the ventral and dorsal sides of each specimen to compensate for the loss of signal quality, due to light scattering, in deeper optical slices, resulting in two views of the same sample.

#### Live-cell imaging (2D)

##### Tissue collection and preparation

HeLa cells expressing endogenously tagged Halo-TRFl were labelled with Janelia Fluor 646 Halo ligand (Promega) in order to mark telomeres. To visualize TERRA, ectopically expressed 15q-TERRA species were tagged with PP7 stem-loop structures that were bound by phage coat protein fused to GFP (PCP-GFP). Live cells were imaged using a Nikon Confocal Spinning Disk microscope equipped with two Photometries Prime 95B cameras and sCMOS Grayscale Chips. Imaging was performed with a 100x objective in an equilibrated incubation chamber at 37°C and 5% CO2. Images were acquired as multi-channel single planes at a rate of 20 frames per second (50 ms exposure, 200 frames per movie).

#### Live-cell imaging (3D)

##### Tissue collection and preparation

COS7 cells were cultured in Dulbecco’s modified Eagle’s mediume (DMEM) supplemented with 10% fetal bovine serum and were incubated at 37°C with 5% CO2. Cells were patterned wither by following either [46] or by light-induced fibrinogen printing controlled by a PRIMO micropatterning machine (Alvéole, France) mounted on an inverted microscope (Olympus 1×81, Japan). Time-lapse imaging of patterned cells was performed by using a 3D Cell Explorer-fluo microscope (Nanolive, Switzerland) with 2 second time interval and a voxel size of (360, 200, 200) nm (Z,Y,X) in phenol-red-free DMEM at 37°C with 5% CO2.

#### Experimental data processing

##### Imaging-based spatial transcriptomics

###### Projection, stitching and alignment

To yield 2D images the acquired stacks were reduced using either maximum intensity projection (MIP) or a custom implementation of extended depth-of-field (EDF, [47]). Tiled acquisitions were stitched together into a mosaic image with Ashlar [48], which uses a variant of phase correlation [49] to compute the offset between the different tiles at subpixel precision [50] in a simultaneous fashion. Only the DAPI channel was used to retrieve the stitching coordinates, and different cycles in the same experiment were stitched independently. After obtaining the mosaics for all cycles in an experiment, we registered them with *wsireg* [51], which uses *elastix* [52, 53] as a backend. We allow for rigid-body alignment as well as non-linear warping, which we found did not aggressively deform the sample and was able to align fine details properly. We only used the DAPI channel for inter-cycle registration.

###### Spot detection (spotiflow)

All results with Spotiflow were obtained using a Spotiflow network trained on the HybISS dataset (*cf*. Fig. 1b, Fig. 1c, Supp. Fig. 6, Supp. Table 2, Supp. Table 3) to detect spots independently across cycles and channels. The probability threshold used was optimized from the validation data of the *HybISS* dataset.

###### Spot detection (LoG/starfish)

We ran LoG using starfish’s implementation, which is based on scikit-image [24], and used different intensity thresholds including the ‘optimal’ one (0.138) which was computed from the training data of the *HybISS* dataset. To ensure a fair comparison, we detected transcripts independently across cycles and channels as done for Spotiflow.

###### Gene decoding

To extract gene expression maps the detected transcripts were assigned a gene using starfish’s implementation of an intensity-based nearest-neighbor decoder. When decoding spots detected via Spotiflow, we fed the decoder the spot probabilities output by the network instead of their raw intensity. Gene expression heatmaps were obtained by performing Gaussian kernel density estimation (KDE) with variance *σ*^2^ = 5 on the gene signal. The heatmaps were clipped to (0,1) after applying percentile-based normalization with *p*_*min*_ = 0, *P*_*max*_ = 99.9.

###### Time/memory benchmarking

To compare the time and memory efficiency of starfish and Spotiflow in an end-to-end setting (*cf*. Fig. 2e, Fig. 2f), we used Unix’s `time` module. We detected spots on an E12.5 mouse embryonic brain using LoG on the maximum intensity projection of the input along the cycle and channel dimensions, as done in previous studies [9, 54]. The spot intensities were then traced back along the non-projected input to retrieve the intensity of spots at the different cycles and channels. For Spotiflow, the detections were done independently on each cycle and channel as it is computationally affordable for larger image sizes.

###### Zero-shot autofluorescence removal

We first built a dataset from a single experiment consisting of a regular *HybISS* acquisition preceded by imaging an only-autofluorescence cycle. After registering both images, we then detected spots (using the Spotiflow model trained on the *HybISS* dataset) in one channel (corresponding to 750 nm) of the autofluorescence cycle and the same channel of the first HybISS cycle. We generated the non-autofluorescent ground-truth by subtracting all the autofluorescent detections that matched (at spatial cutoff *c* = 3) to a detection in the HybISS channel. We generated three spatially-disjoint splits from this experiment (training, validation and test). We finally fine-tuned the Spotiflow model pre-trained on *HybISS* on the generated training dataset to predict the non-autofluorescent spots. Quantification is reported on the test split (*cf*. Fig. 3d,e).

###### Views fusion

Two views were combined using a three-step registration procedure, and a composite volume was created based on the registered views. Only the DAPI channel was used for finding the transform, but the transform was applied to all channels.

###### Pre-alignment

During data acquisition, the animal can be positioned arbitrarily. Both images were smoothed using an intensity threshold to obtain a point cloud. Then principal component analysis (PCA) was performed on the point cloud and the image was rotated to align the first principle component with the *x* -axis. After that, an heuristic approach was used to make sure that all animals have the same orientation (with the head of the animal towards 0), based on the fact that the head of the animal has more signal than the rest of the body.

###### Rigid alignment

Pre-aligned images were registered using the Euler transform implemented in *elastix* [52, 53]. First, the registration was done by optimizing the mutual information metric with 5 levels in the image pyramid schedule (64,32,8,4 and 1) and 20000 spatial samples for the metric and gradient estimations. In the second step, rigid registration was done by optimizing the normalized correlation coefficient (NCC) using 5 levels in the image pyramid schedule (16, 8, 4, 2 and 1) and 50000 spatial samples.

###### Deformable alignment

Due to small deformations resulting from the sample handling between the acquisition of the views, it was not possible to fully register two views using only rigid registration, *elastix’s* B-Spline deformable registration was used as the last registration step. The metrics optimized were the NCC, the mutual information along with a rigidity penalty with the corresponding weights of 1, 1 and 100. Registration was performed in one step at the original resolution with the final grid spacing of 64 voxels in the *Y* and *X* directions and 16 voxels in the *Z* direction using 100000 spatial samples for the metric and gradient estimations.

###### Composite volume

Small inaccuracies in registration of the two views can cause duplication of the spots, therefore instead of directly averaging the registered views, a composite volume was constructed. The DAPI channel of both registered images was smoothed using a Gaussian of kernel size (10, 20, 20) (ZYX). The volume was then split into the areas where one of the smoothed views had higher intensity than the other, and this mask was used to create a weighted average of the views.

###### Spot detection with Spotiflow

The Spotiflow model used was pre-trained on the *synthetic-3D* dataset (with 5 = 2) and then fine-tuned for 400 epochs on nine annotated subvolumes and validated on six annotated subvolumes. The spot probability threshold used for the predictions is 0.4.

###### Spot detection with Laplacian-of-Gaussian (LoG)

The *scikit-image* implementation of the LoG blob detector was used with *σ* ∈ [1, 3] and different intensity thresholds (from 0.1 to 0.7 in steps of 0.05). In order to scale to the full volume, we apply the LoG blob detection in a sliding window approach in windows of size (32, 128, 128) (ZYX).

###### Semantic segmentation of DAPI channel

A tissue semantic segmentation mask was obtained by using Otsu’s thresholding on the DAPI channel followed by morphology operations in order to remove small objects (< 100 px.) and small holes (< 100 px.) with an area opening and an area closing respectively. A closing with a cubic structuring element of (3,3,3) (ZYX) was then performed followed by a dilation of a (3,9,9) (ZYX) cuboid in order to expand the mask a few voxels from the stained nuclei.

###### Time-lapse processing with Spotiflow

Resulting time-lapses were first affinely aligned on Fiji [55] (using the *descriptor-based registration (2d/3d*) [56] plugin) to an image with fluorescently labelled beads as a reference. Movies were first spatially cropped so that each crop contains only one cell. In each crop, spots were detected independently per frame and channel, where one channel contains the Telomeres marker and the other the TERRA marker, using the deepBlink and Spotiflow models trained on the *Telomeres* and *Terra* datasets, which consist of annotated frames of the movies obtained by the procedure described above. The optimized probability threshold on each dataset was used for Spotiflow. For deepBlink, the probability threshold was set to the default (0.5). Single particle tracking was performed using Trackmate [25] with a spot radius of 0.15 *µm* and simple LAP tracking with the following parameters for Telomeres/TERRA: linking maximum distance 0.22/0.60 *µm*, gap-closing maximum distance 0.44/1.0 *µm* and gap-closing maximum frame gap 10 frames.

###### Time-lapse processing with Spotiflow

In order to fine-tune the Spotiflow model pre-trained on the dataset *synthetic-3D*, the first two frames of one of the volumetric movies were annotated using *napari*. The Spotiflow network was then fine-tuned for 10 epochs on the two frames, which were treated as independent 3D stacks. The fine-tuned Spotiflow model was afterwards used to detect spots (corresponding to the lipid droplets) in an out-of-training movie across the whole duration (200 frames). The localizations were tracked across time using the Crocker-Grier algorithm [57] implemented in *Trackpy* [30], using a gap closing of 2 and a search radius of 5 px. Tracks shorter than 20 frames were discarded.

###### Spot detection with Spotiflow

The Spotiflow model pre-trained on the dataset *synthetic-3D* and fine-tuned on the 3D smFISH *Platynereis dumerilii* annotated subvolumes (see above) was used to detect spots on a 159GB EASI-FISH volume of a lateral hypothalamus section of a mouse brain [13].

###### Distributed inference

To speedup inference, we wrote a custom parallelization procedure which distributes overlapping tiles across different GPUs. While overlapping tiles are widely used to run CNNs on non GPU memory-fitting inputs in order to avoid border artifacts, we also exploit the strategy to distribute these tiles across different GPUs, achieving quasi-linear speedup w.r.t. the number of GPUs. The procedure is based on the tile iterator implemented in the Python package *csbdeep* [58], which we wrap as a *torch.data.utils.Dataset* instance as well as includes a custom dataloading sampler adapted from [59]. As data needs to be lazily loaded due to its large size, the ability of *torch.data.utils.DataLoader* to spawn several workers allows loading the next tiles (*pre-fetching*) asynchronously, greatly reducing I/O bottlenecks. After the workload of each GPU is finished, shapes of the tensors containing spots are sent to the process of the main GPU, which uses them to generate appopriate-sized empty tensors which will finally be filled by gathering the spot results of each GPU. Distributed prediction can be run using the *torch.distributed* protocol seamlessly in a system where the amount of GPUs is greater than 1 using *torchrun*.

