## Supplementary material for "Spotiflow: accurate and efficient spot detection for fluorescence microscopy with deep stereographic flow regression": Supp Material

**4 References** **24**

### 1 Supplementary Figures

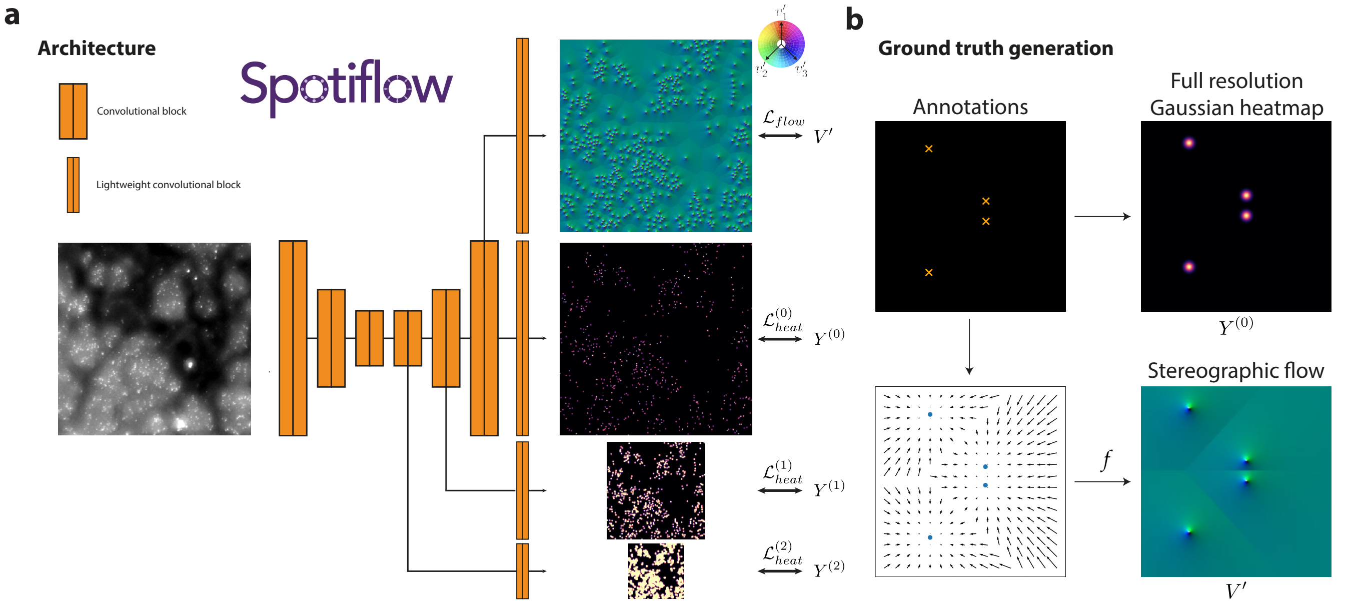

**Supplementary Figure 1. Architecture of Spotiflow. a)** An input is processed by a U-Net, which consists of *convolutional blocks* of depth $d = 3$ and increasing feature maps (16, 32, 64 channels) and *lightweight convolutional blocks* of depth $d = 3$ and non-increasing feature maps (1 channel). All convolution kernels are 3×3 (3×3×3 in the 3D case). The combined loss is the summ of $L$ multiscale heatmap losses $\mathcal{L}^{(i)}_{heat}$ (binary cross-entropy loss) and the stereographic flow loss $\mathcal{L}_{flow}$ ($L_1$ loss). **b)** Ground truth generation from point annotations for training Spotiflow. First, a Gaussian is generated on top of every annotation yielding the full resolution Gaussian heatmap $Y^{(0)}$. This heatmap is then further processed to obtain its representation at different resolutions (*multiscale heatmaps*). Second, a *local offset vector field* is built in which every position contains the vector directed to the closest ground truth spot. The stereographic flow is then obtained by computing the inverse stereographic projection position-wise.

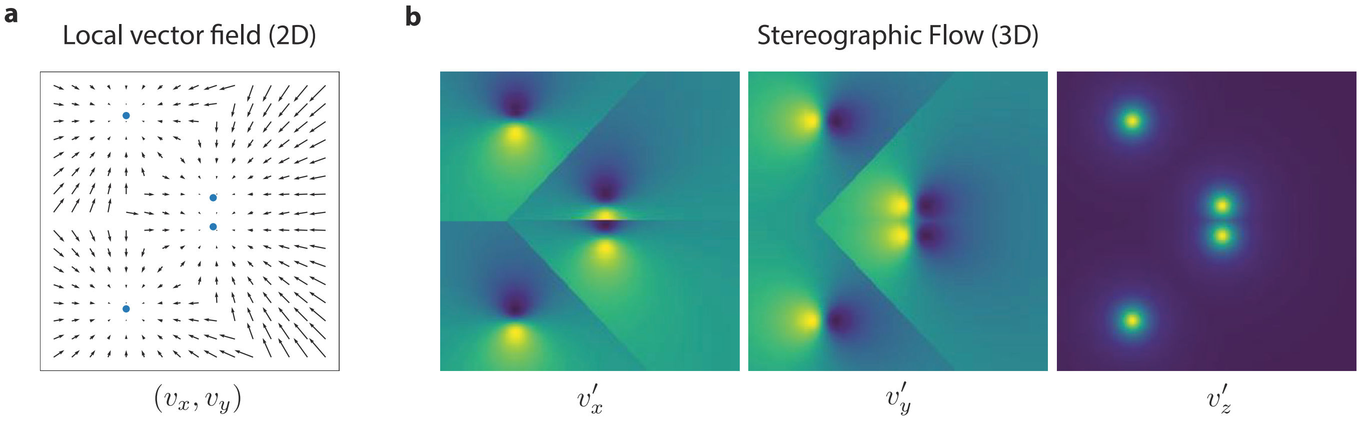

**Supplementary Figure 2. Principle of stereographic flow (2D). a)** For given ground truth center locations (blue dots) the 2D vector field $\{(v_x, v_y)_{ij}\}$ is defined as the vector from each pixel $ij$ to the nearest ground truth spot. **b)** The 2D vector field is embedded in $\mathbb{R}^3$ onto the unit 3-dimensional sphere $S^2$ via an inverse stereographic projection yielding the 3D *stereographic flow* $(v'_x, v'_y, v'_z)$.

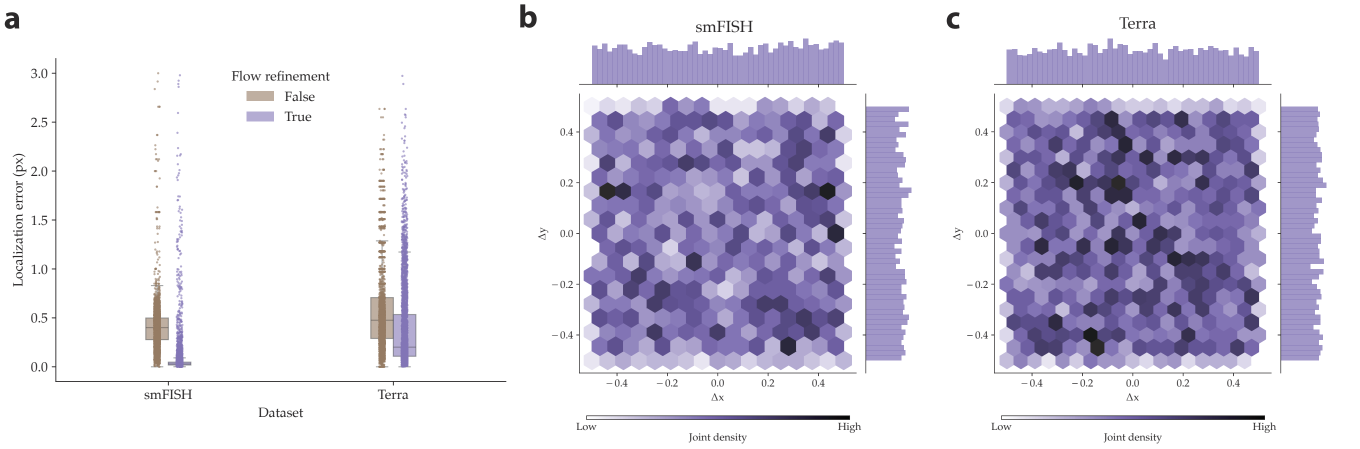

**Supplementary Figure 3. Assessment of Spotiflow's subpixel refinement via stereographic flow. a)** Localization error for true positive detections on all test images for *smFISH* and *Terra* with and without flow refinement. **b)** Joint distribution of the 2D spatial subpixel component $(\Delta x, \Delta y) \in (-0.5, 0.5)^2$ of the prediction of Spotiflow on a test image of the *smFISH* dataset. **c)** Same for an image of the *Terra* dataset. The Spotiflow models used were trained on the respective datasets.

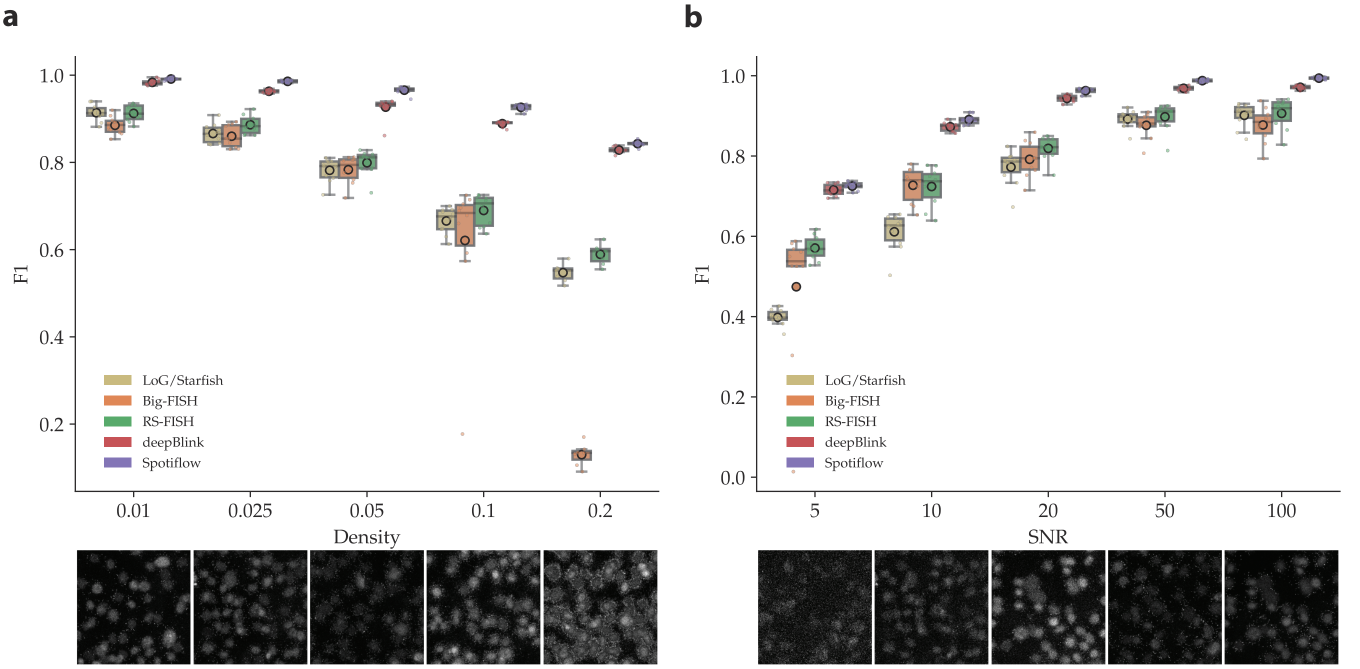

**Supplementary Figure 4. Influence of different experimental conditions on spot detection algorithms. a)** Synthetically generated images with increasing density of spots. Shown is the $F_1$ score with median and interquartile range across all images (boxplot) and individual scores per image (dots), as well as a representative image for each condition. **b)** Same for synthetic data with increasing signal-to-noise ratio (SNR).

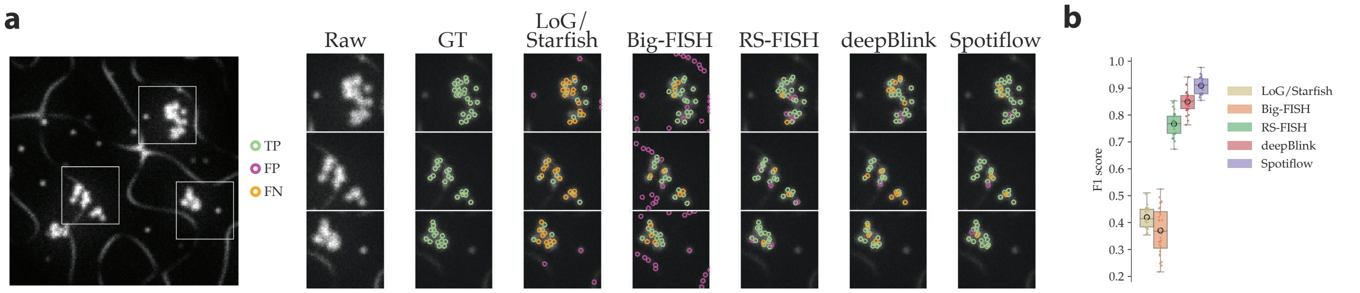

**Supplementary Figure 5. Spot detection in dense clusters. a)** Overview of a test image of synthetic data generated containing spots distributed in constellations. The simulation includes isolated spots, which should not be detected. Depicted in the insets are ground truth and predictions of different methods which were optimized/trained on the training split ($N = 240$) of the dataset. **b)** Benchmarking of methods on the whole test split ($N = 28$) of the dataset. Shown is the $F_1@3$.

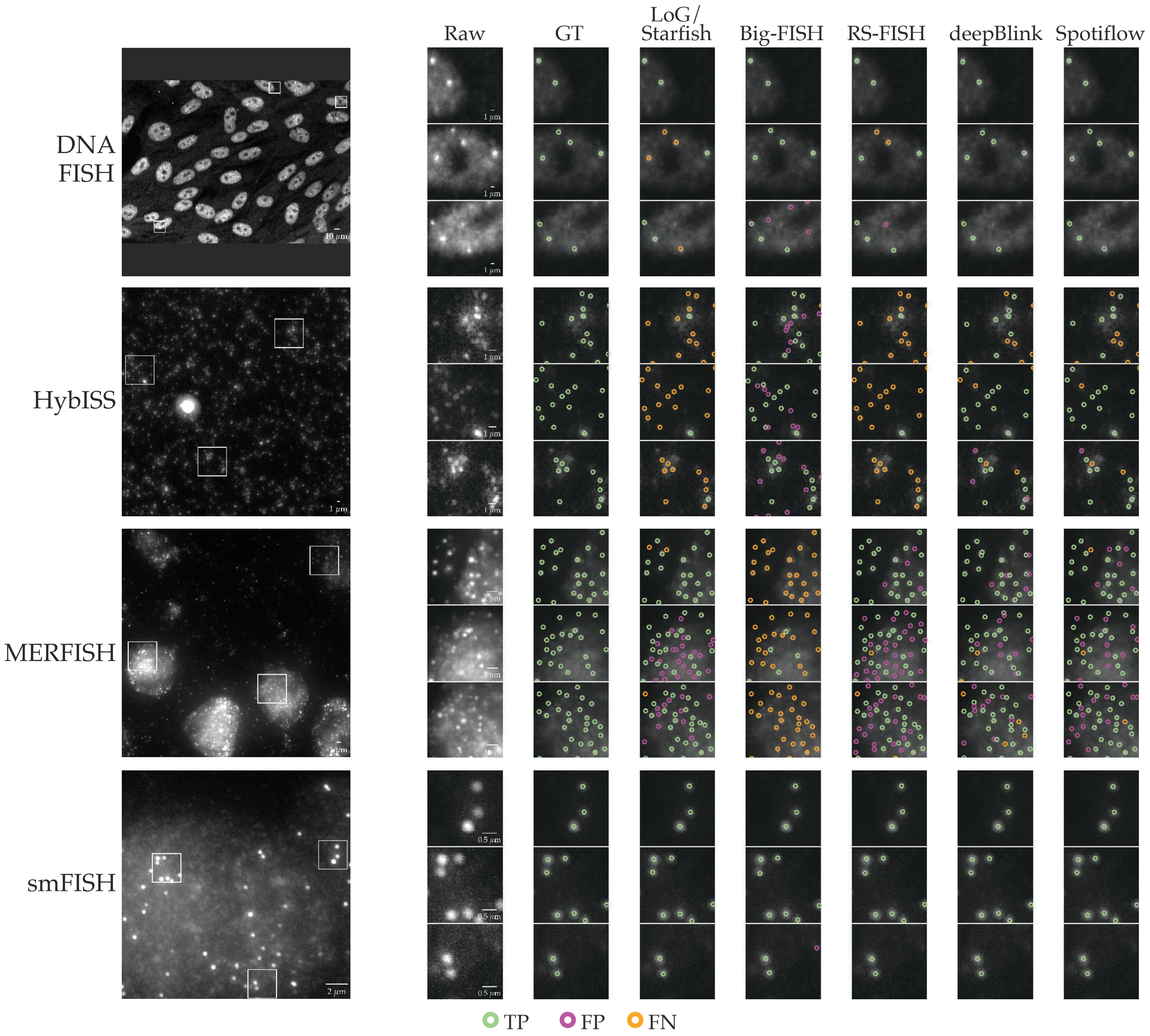

**Supplementary Figure 6. Example images for 2D FISH benchmark datasets and corresponding spot predictions of compared detection methods.** $TP$, $FP$ and $FN$ denote true positive, false positive and false negative detections respectively.

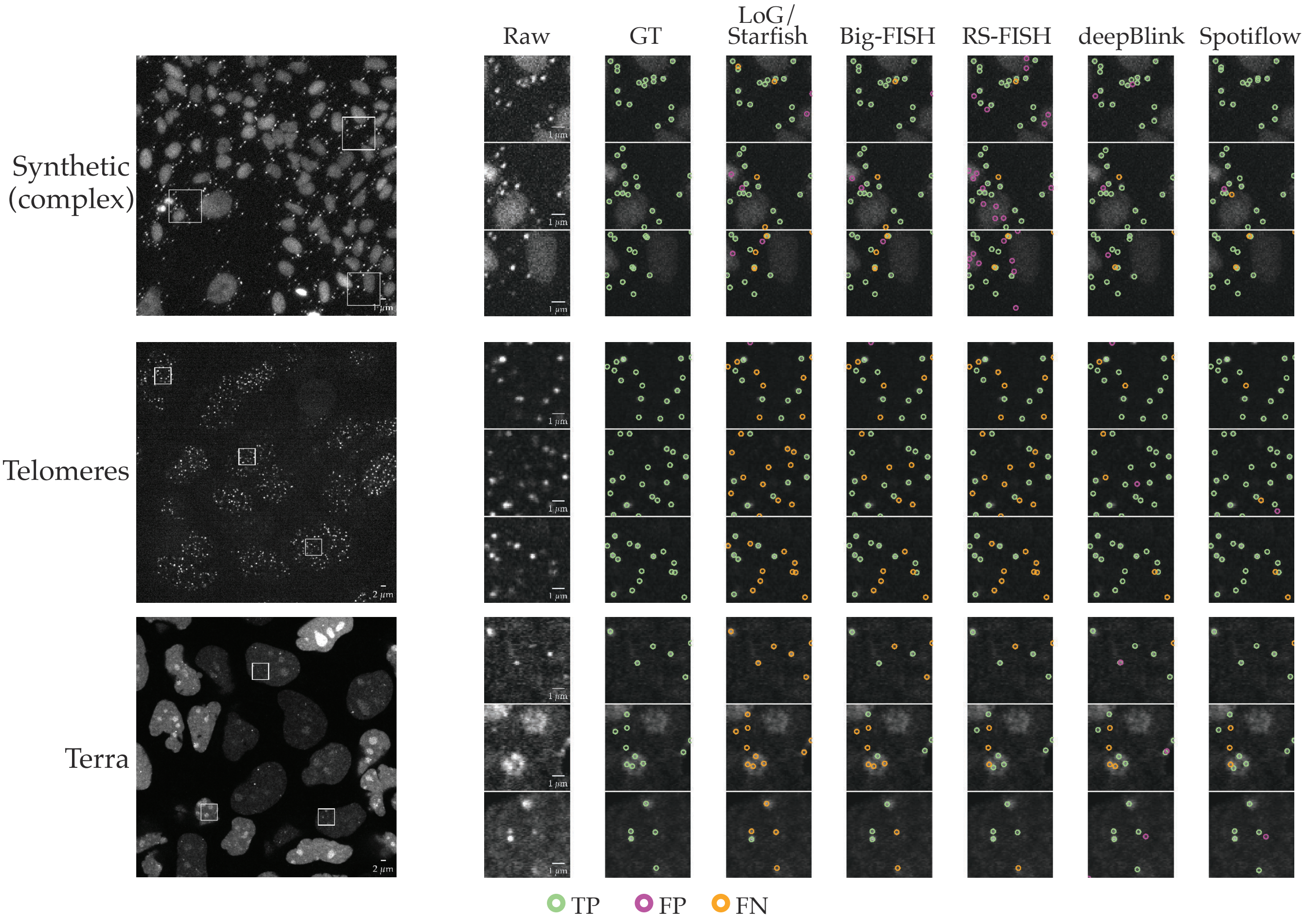

**Supplementary Figure 7. Example images for 2D synthetic and live-cell benchmark datasets and corresponding spot predictions of compared detection methods.** $TP$, $FP$ and $FN$ denote true positive, false positive and false negative detections respectively.

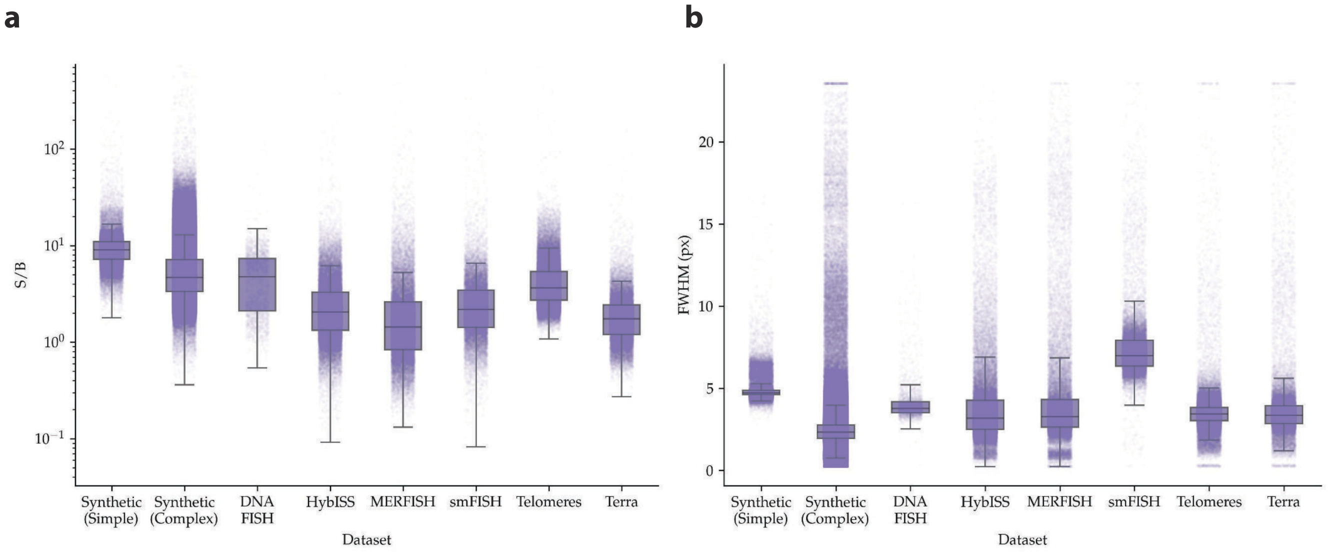

**Supplementary Figure 8. Estimated signal-to-background ratio (S/B) and full width at half maximum (FWHM) of ground truth spots in all benchmark datasets.** S/B (**a**) and FWHM (**b**) are estimated by fitting Gaussian functions in a small window centered around a ground truth spot.

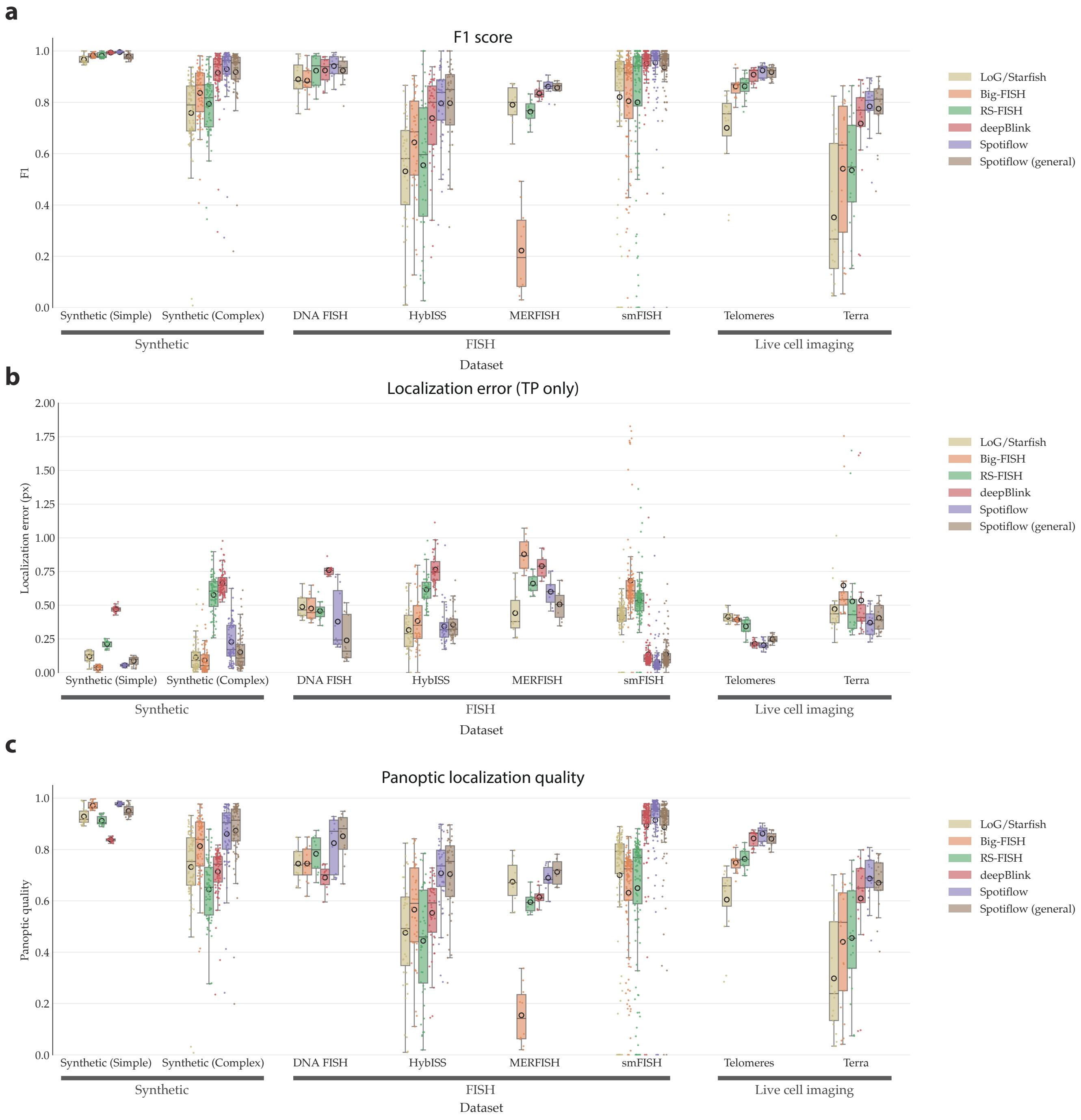

**Supplementary Figure 9. Evaluation of 2D spot detection methods on benchmark datasets.** Shown are **a)** F1-score (higher is better), **b)** Localization error in pixels (lower is better) and **c)** Panoptic Quality score (higher is better) for different methods on different synthetic and real datasets. Compared methods are: Starfish [1], Big-FISH [2], RS-FISH [3], and deepBlink [4]. Shown are median and interquartile ranges across all images (boxplot) and individual scores per image (dots).

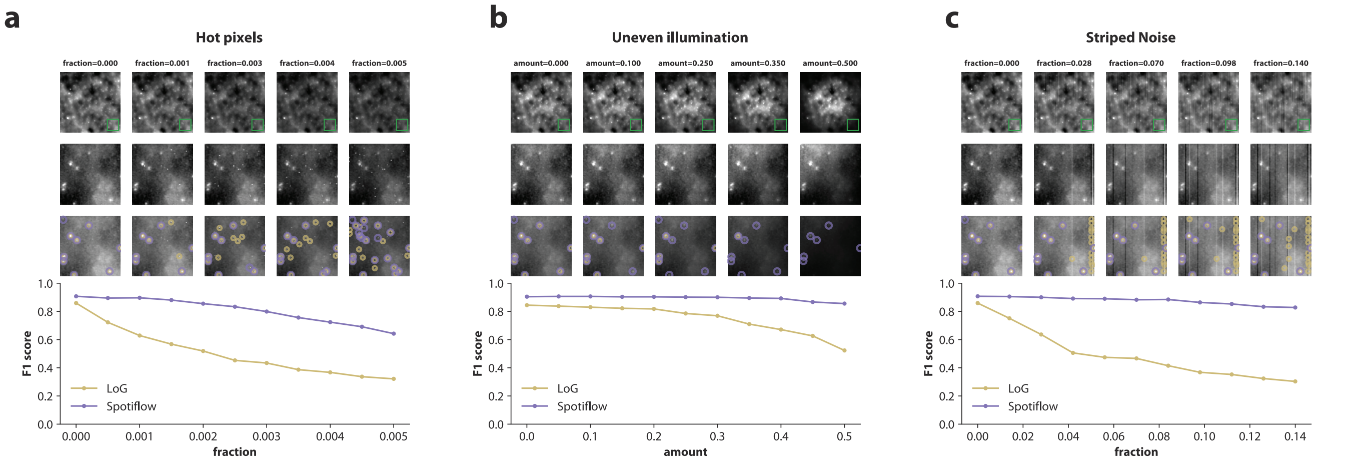

**Supplementary Figure 10. Assessment of the impact of common image artifacts on spot detection performance.** For a test image of the *HybISS* dataset an increasing amount of different types of artifacts were introduced. Shown is the $F_1$ score for Spotiflow and LoG. **a)** Hot pixels, commonly present in images obtained from CCD cameras. **b)** Vignetting effects caused by uneven illumination. **c)** Vertical-striped noise from CCD cameras. Results show the F1 score (higher is better) for different levels.

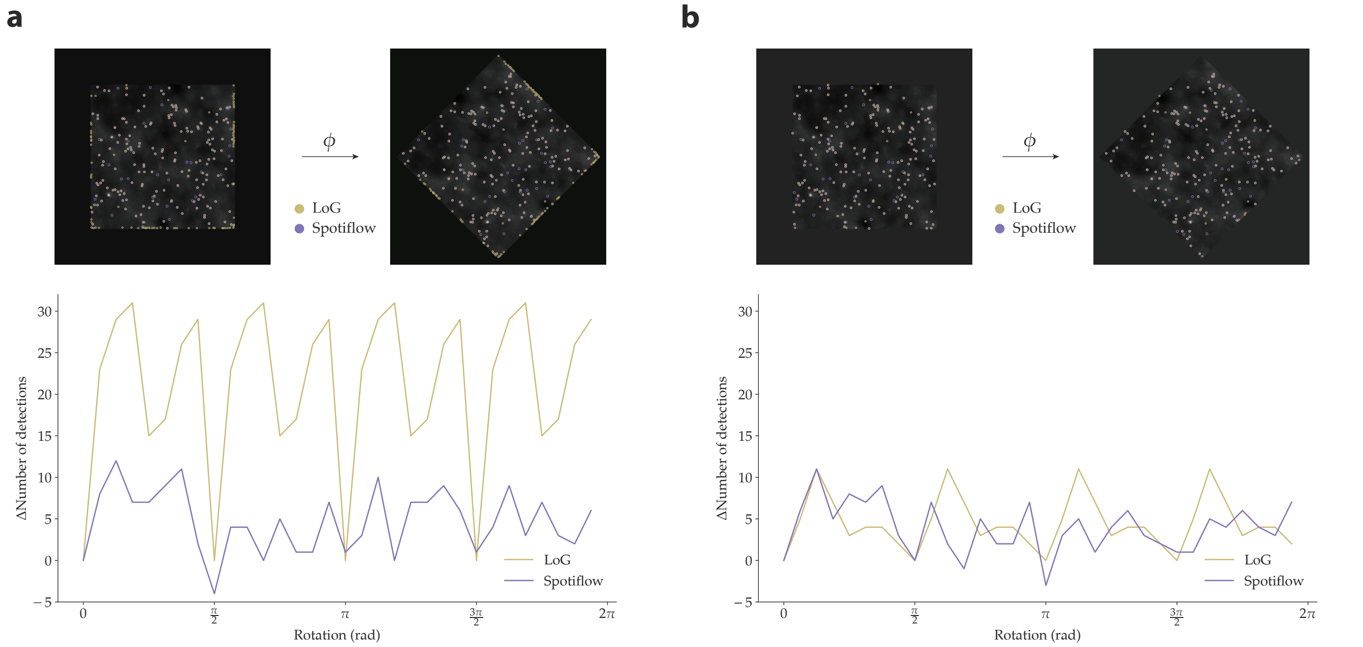

**Supplementary Figure 11. Effect of input rotation on spot detection with LoG and Spotiflow.** LoG (optimal threshold from *HybISS* training data) and Spotiflow (*HybISS* model) are run on an image from the *HybISS* dataset which is rotated at varying angles $\varphi$ with cubic spline interpolation. The impact of rotation is assessed with the difference to the number of detections obtained with the original (non-rotated) input. Results with different rotation padding strategies, zero padding (**a**)) and constant padding with the average intensity at the border of the image (**b**)), are shown.

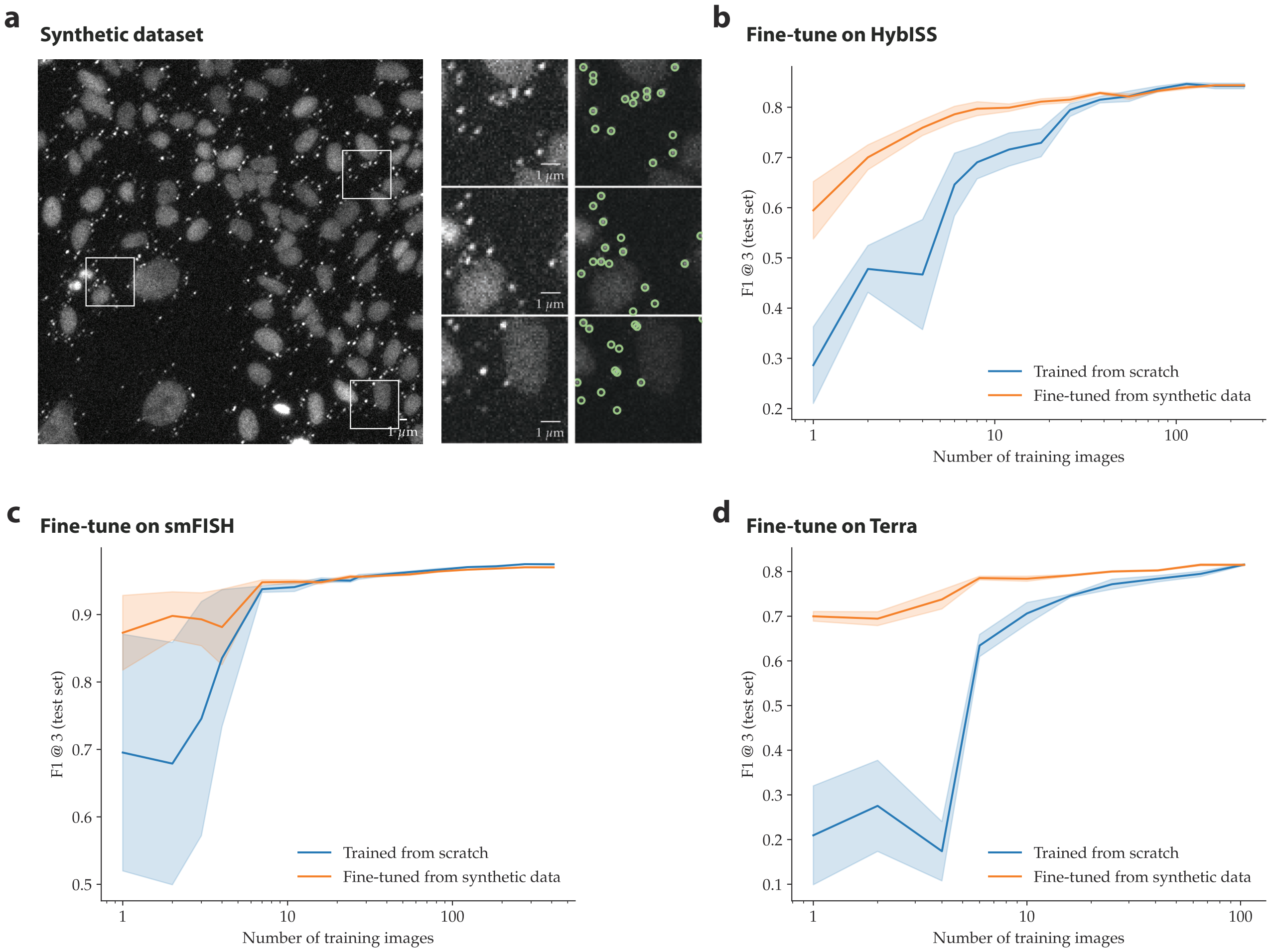

**Supplementary Figure 12. Fine-tuning Spotiflow from synthetic data.** **a)** We fine-tune a Spotiflow network pre-trained on a realistically-simulated synthetic dataset on several datasets: **b)** on HybISS (iST), **c)** on smFISH (iST), and **d)** on Terra (live-cell). Line plots show the F1 score (higher is better) on the whole test split of each dataset after fine-tuning on subsets of different size of the training split.

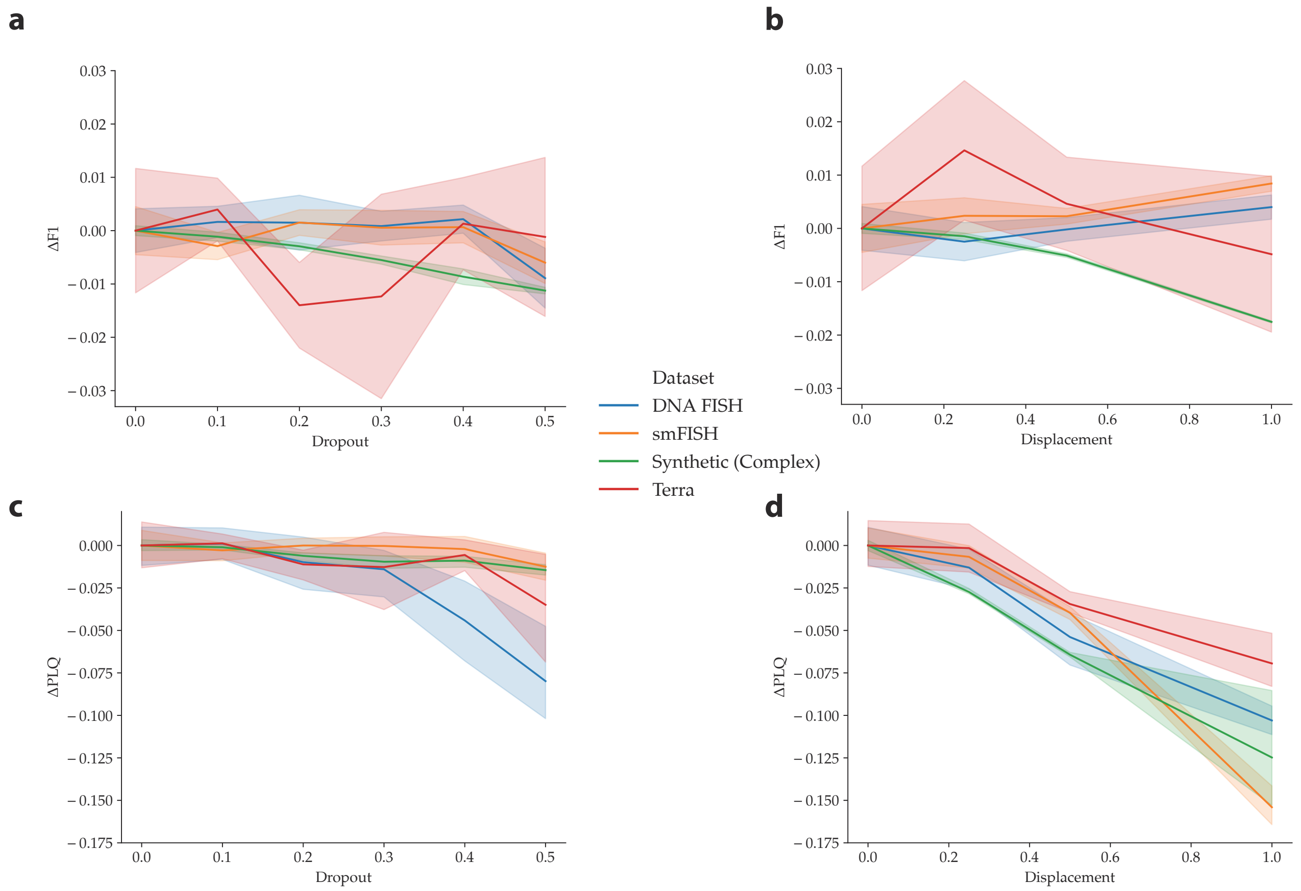

**Supplementary Figure 13. Robustness assessment of Spotiflow to annotation quality.** Training ground truth is perturbed by either dropping out a fraction of annotations (**a)** and **c)**) from each training image or by randomly displacing the annotation around the spot center (**b)**, **d)**). The difference in $F_1@3$ ($\Delta F_1$), to the baseline (no perturbations) is shown in **a)** and **b)**, while the difference in $PLQ$ ($\Delta PLQ$) is depicted in **c)** and **d)**. The evaluation is performed on the original unperturbed test split of the dataset.

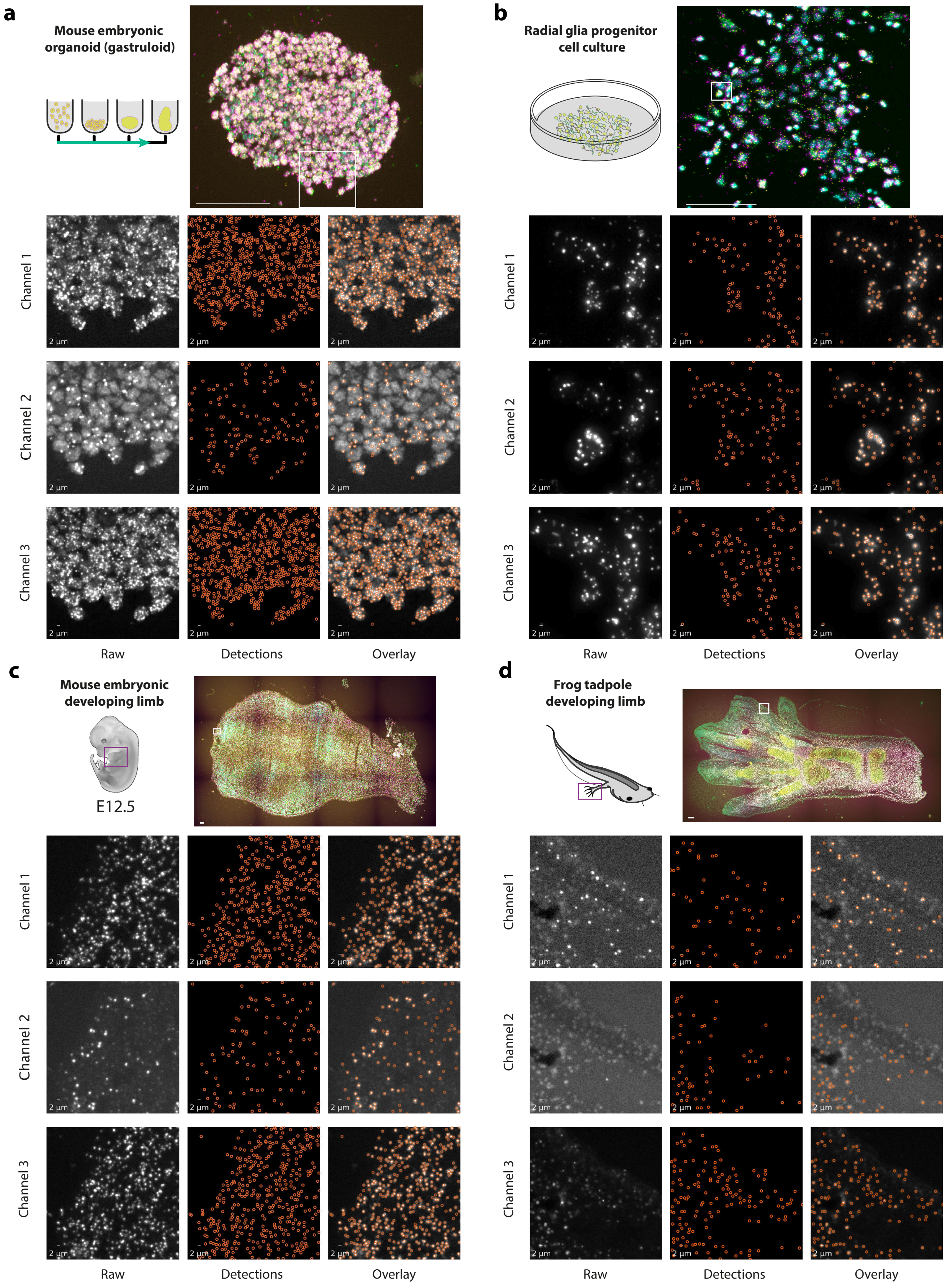

**Supplementary Figure 14. Spot detection results of Spotiflow for different HybISS samples. a)** Mouse gastruloid. **b)** Radial glia progenitor (RGP) cells culture. **c)** Mouse embryo developing limb. **d)** *Xenopus laevis* developing limb. The Spotiflow model used was trained on *HybISS*, which contains only mouse embryo data, and was run at the optimal probability threshold as optimized during the training stage. Scalebars in the overview images correspond to 100$\mu$m.

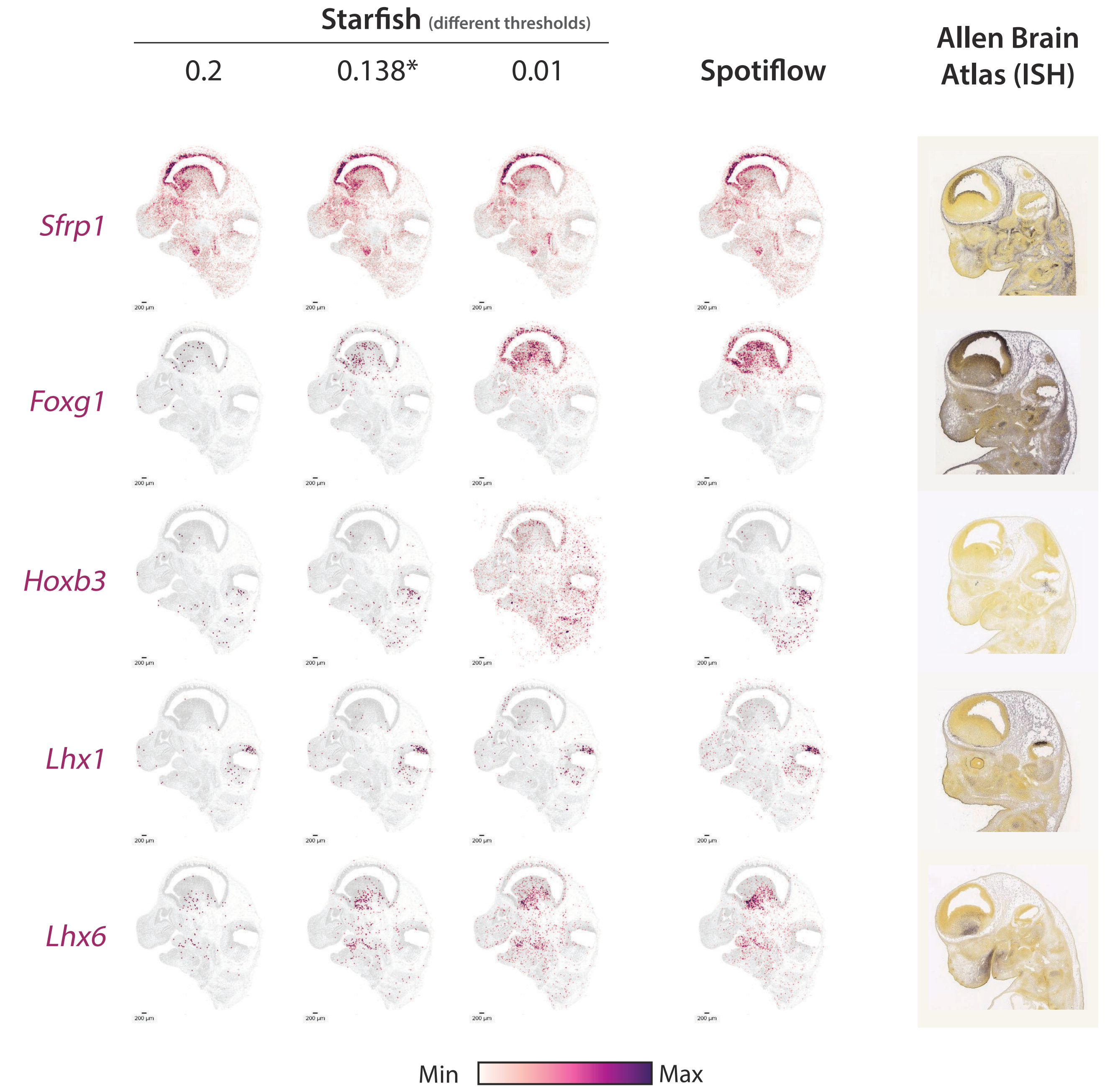

**Supplementary Figure 15. Comparison of HybISS gene expression maps obtained using Starfish and Spotiflow for an E13.5 embryonic mouse brain.** Starfish is run at different intensity thresholds, including the optimal one according to the training split of the *HybISS* dataset (0.138). The Spotiflow model used was trained on *HybISS* and is run at the optimal probability threshold as optimized during training. Allen Brain Atlas (ISH) for similar slides is shown as a reference.

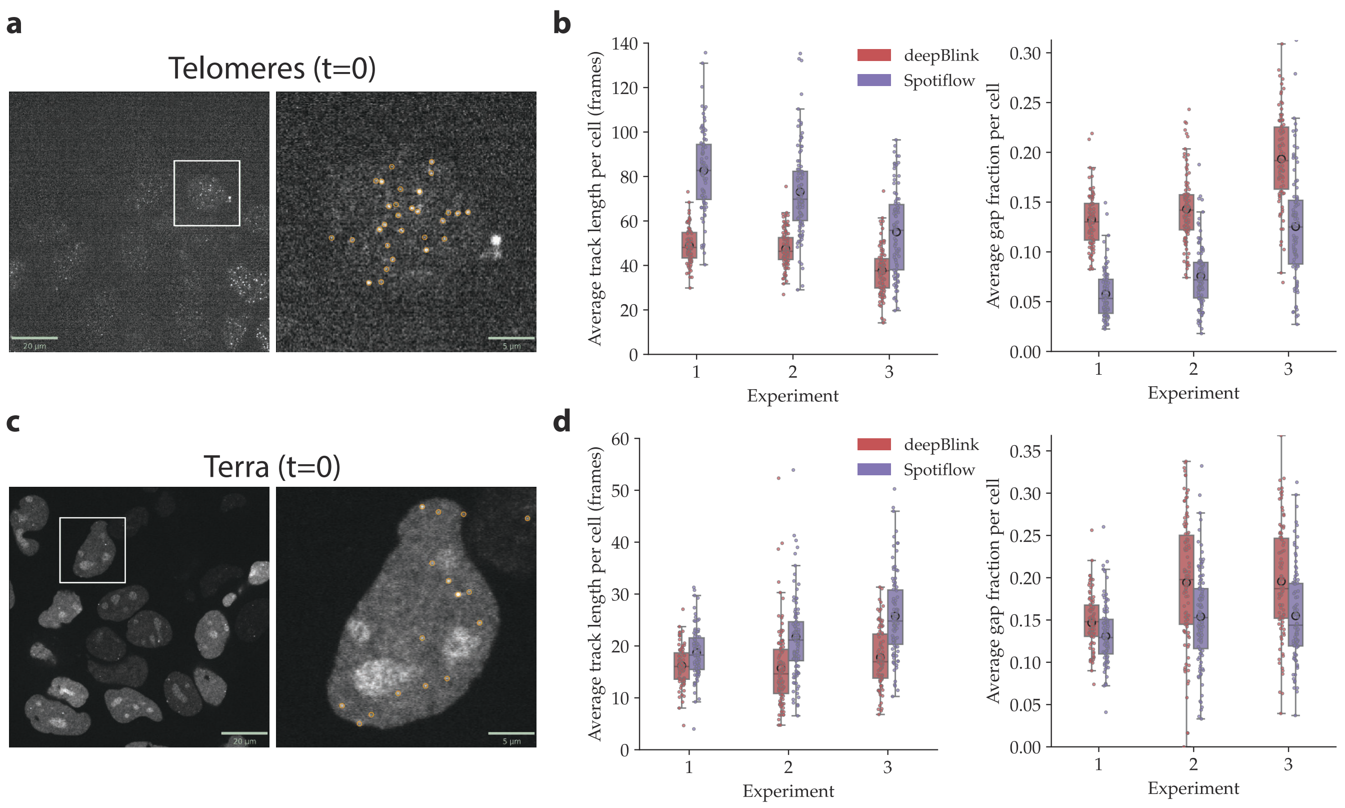

**Supplementary Figure 16. Tracking of spots in live-cell microscopy movies. a)** Single frame of a live-cell aquisition of HeLa cells with labeled Telomeres (orange). **b)** Average length of tracks per cell after spot-detection and tracking via TrackMate [5]. Shown are tracklength distributions for three different experimental conditions. **c)** Single frame of a live-cell acquisition of HeLa cells with labeled Terra (orange). **d)** Average length of tracks per cell as in **b**.

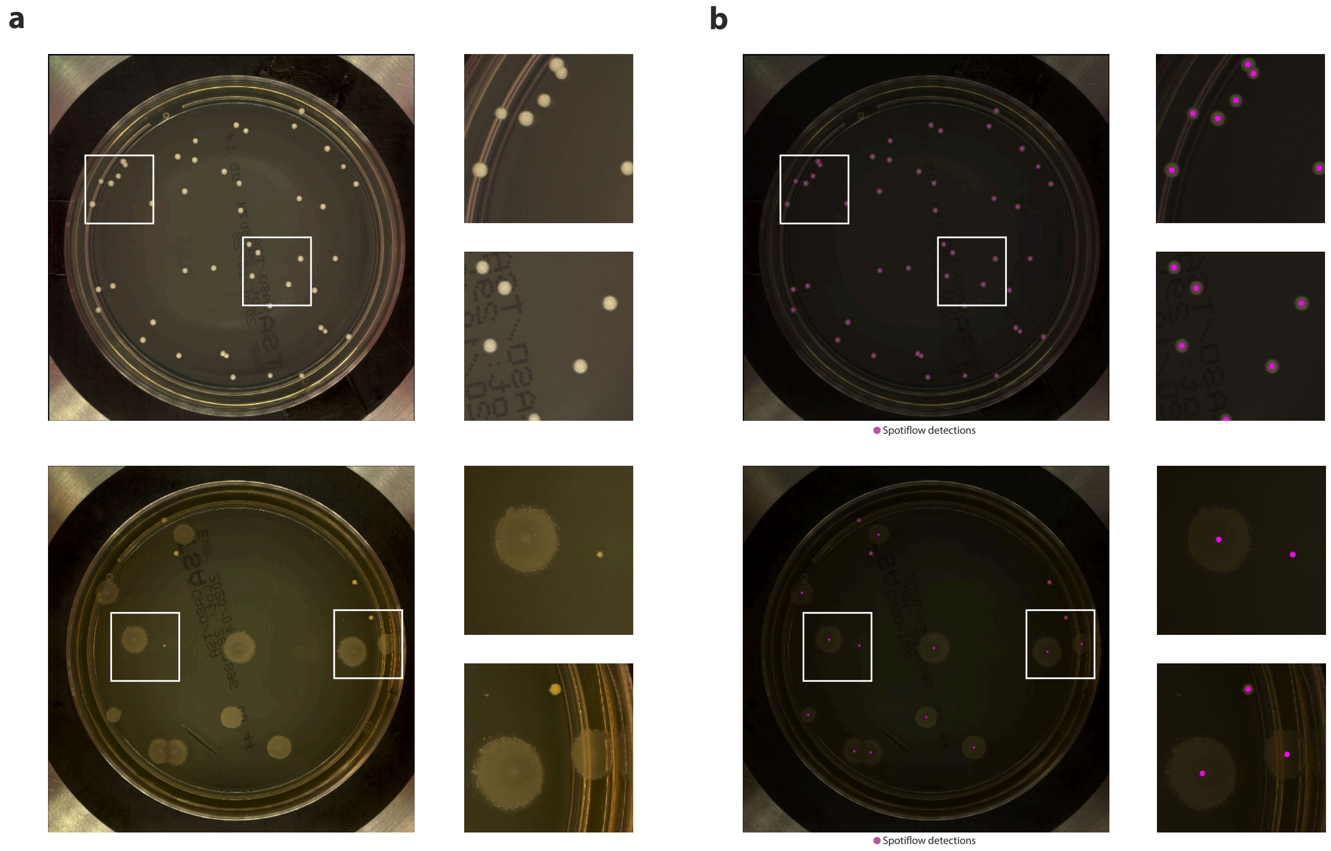

**Supplementary Figure 17. Application of Spotiflow as a general object detection method: bacteria colony detection.** Use case of Spotiflow's capability as a general detection method for detecting bacteria colonies. Spotiflow was trained on RGB images from the *AGAR* dataset [6] ($N = 3152$). **a)** Raw RGB images from the test split ($N = 1106$) of the dataset. **b)** Overlaid detections of Spotiflow on the same test images.

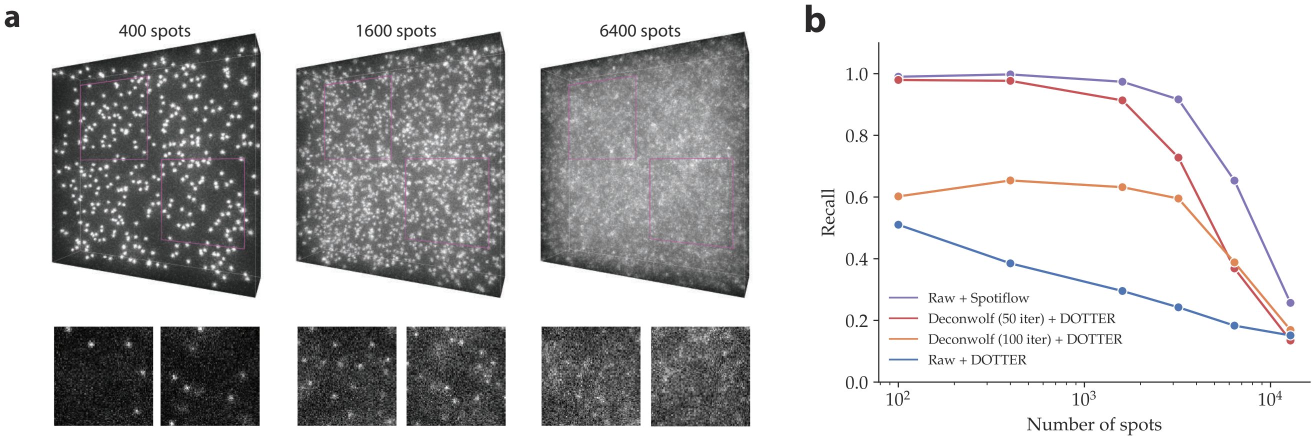

**Supplementary Figure 18. Assessment of Spotiflow performance compared to deconvolution-aided spot detection. a)** Overview of raw synthetic volumes simulated with different densities (400, 1600 and 6400 spots per volume respectively). Data from [7]. **b)** Performance assessment of Spotiflow and the 0th (raw), 50th and 100th last iteration of the deconvolution tool *Deconwolf* followed by an intensity-based spot detection method (DOTTER) [7] along with a Spotiflow model trained on *synthetic-3D* (*cf.* Methods). Percentage of ground truth spots detected (recall) is shown on the y-axis, different number of spot per volume on the x-axis.

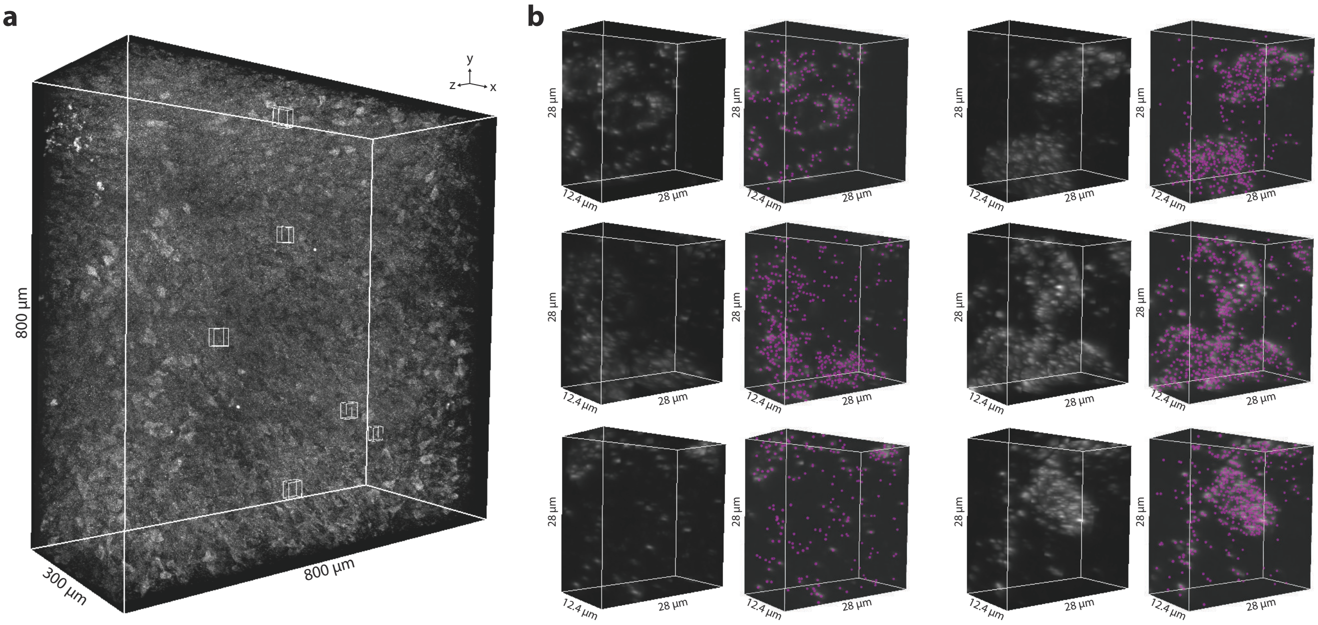

**Supplementary Figure 19. Spotiflow predictions on EASI-FISH data. a)** Overview of a lateral hypothalamus section of a mouse brain processed using EASI-FISH. Data from [8]. **b)** Pairs of insets of raw data (left) and with overlaid detections of Spotiflow (right) of the same stack.

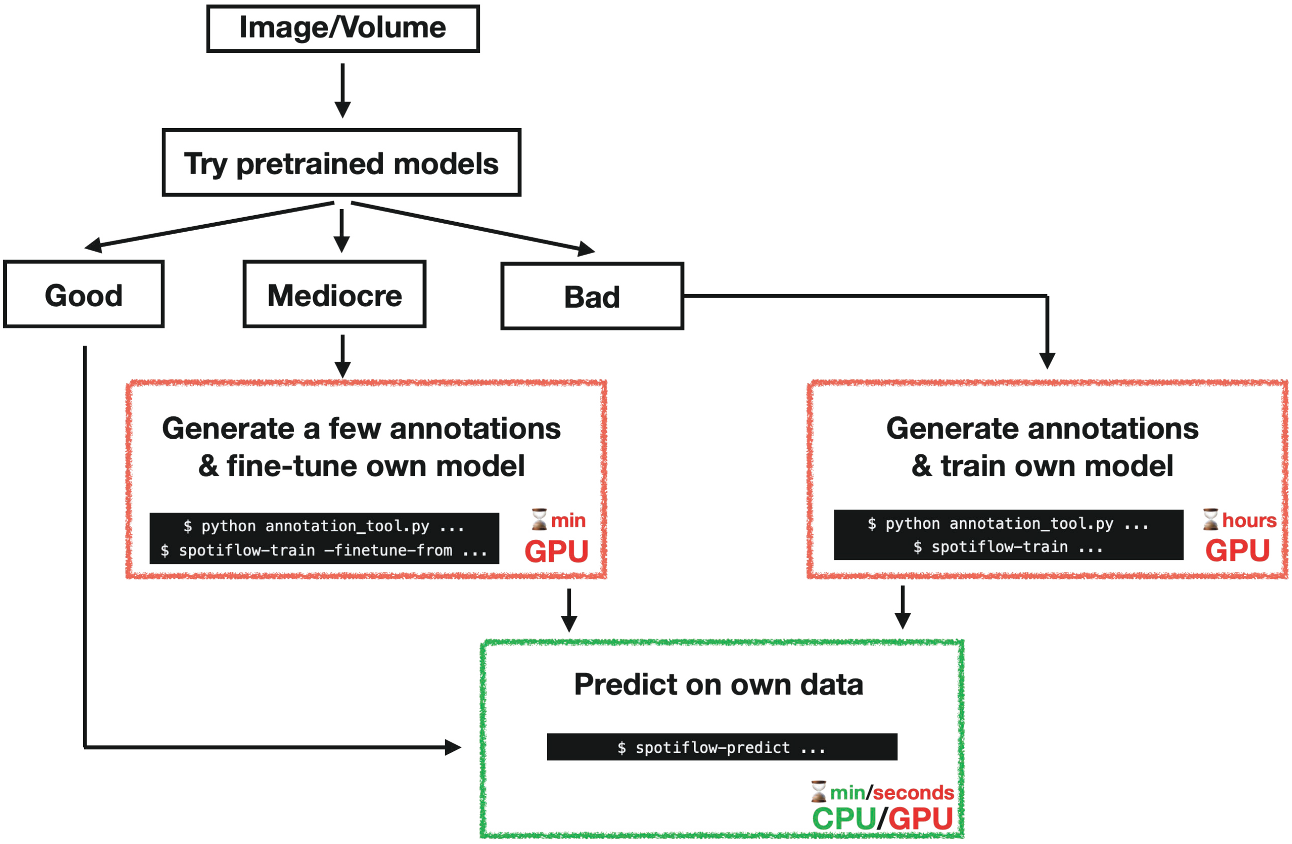

**Supplementary Figure 20. Flowchart depicting the adoption procedure of Spotiflow for a new user.** The time estimates refer to the average computational workload expected for that step for middle-sized images.

#### 2 Supplementary Tables

| Dataset | Multiscale loss | Stereographic flow | $F_1$ | $PLQ$ |
|---|---|---|---|---|
| smFISH | - | - | 0.955±0.005 | 0.813±0.004 |
| | ✓ | - | 0.957±0.005 | 0.814±0.005 |
| | - | ✓ | 0.958±0.005 | **0.916±0.006** |
| | ✓ | ✓ | **0.959±0.005** | **0.916±0.007** |
| Terra | - | - | 0.785±0.003 | 0.639±0.003 |
| | ✓ | - | 0.791±0.002 | 0.644±0.003 |
| | - | ✓ | 0.786±0.004 | 0.684±0.004 |
| | ✓ | ✓ | **0.794±0.005** | **0.694±0.003** |

**Supplementary Table 1. Ablation study for the multiscale loss and the stereographic vector field regression.** Metrics were aggregated over 5 different runs per combination.

| Dataset | Source | Modality | Subpixel annotations | #Train | #Validation | #Test | #Spots/img | Width/Height (px) | Pixel size ($\mu$m) | FWHM (px) | S/B |
|---|---|---|---|---|---|---|---|---|---|---|---|
| Synthetic (Simple) | – | Synthetic | – | 100 | 20 | 20 | 349±164 | 512 | - | 4.69±0.12 | 8.48±2.02 |
| Synthetic (Complex) | – | Synthetic | – | 300 | 20 | 80 | 1100±1096 | 512 | 0.10 | 2.28±0.38 | 4.23±1.42 |
| DNA FISH | [9] | FISH | ✓ | 21 | 3 | 12 | 78±30 | 930, 1306 | 0.32 | 3.77±0.32 | 4.89±2.73 |
| HybISS | – | FISH | – | 137 | 39 | 44 | 221±369 | 512, 1024 | 0.15, 0.32, 0.34 | 3.07±0.63 | 1.88±0.75 |
| MERFISH | [10] | FISH | – | 12 | 6 | 10 | 1234±225 | 512 | 0.11 | 3.19±0.65 | 1.31±0.65 |
| smFISH | [4, 11] | FISH | ✓ | 411 | 103 | 129 | 39±38 | 512 | 0.04 | 6.97±0.75 | 2.19±0.93 |
| Telomeres | – | Live-cell | ✓ | 70 | 15 | 15 | 612±262 | 1024 | 0.11 | 3.43±0.40 | 3.64±1.13 |
| Terra | – | Live-cell | ✓ | 104 | 23 | 21 | 218±129 | 1024 | 0.11 | 3.34±0.54 | 1.74±0.60 |

**Supplementary Table 2. Descriptive summary of benchmarking datasets.** The dash symbol (–) in the source column indicates that the dataset has been obtained internally.

| Dataset | Modality | Manufacturer | NA | Objective magnification | Fluorophore(s) |
|---|---|---|---|---|---|
| DNA FISH | Confocal Spinning Disk | PerkinElmer | 0.9 | 40X | Cy5, AF568, AF488 |
| | Epifluorescence | Leica | 0.75 | 20X | AF750, Cy3, Cy5, AF488 |
| HybISS | Epifluorescence | Nikon | 0.8 | 20X | AF750, Cy3, Cy5, AF488 |
| | Epifluorescence | Zeiss | 0.8 | 20X | AF750, Cy3, Cy5, AF488 |
| MERFISH[10] | Epifluorescence | Nikon (+ custom) | 0.95 | 60X | AF750, Cy5 |
| smFISH[4, 11] | Epifluorescence | Zeiss | 1.4 | 100X | Quasar670, Quasar570 |
| Telomeres | Confocal Spinning Disk | Nikon | 1.49 | 100X | JF646 |
| Terra | Confocal Spinning Disk | Nikon | 1.49 | 100X | GFP |

**Supplementary Table 3. Specification of optics and fluorophores used to acquire the training datasets.**

| Dataset | Method | F1 | F1-AUC | Accuracy | PLQ |
|---|---|---|---|---|---|
| Synthetic (Complex) | LoG/Starfish | 0.758±0.155 | 0.757±0.155 | 0.631±0.165 | 0.732±0.160 |
| | Big-FISH | 0.836±0.103 | 0.836±0.103 | 0.731±0.140 | 0.813±0.117 |
| | RS-FISH | 0.797±0.119 | 0.796±0.120 | 0.677±0.150 | 0.719±0.123 |
| | deepBlink | 0.915±0.103 | 0.888±0.102 | 0.856±0.135 | 0.713±0.094 |
| | Spotiflow | **0.929±0.105** | **0.927±0.106** | **0.880±0.134** | 0.861±0.123 |
| | Spotiflow (general) | 0.922±0.108 | 0.921±0.109 | 0.869±0.138 | **0.881±0.121** |
| DNA FISH | LoG/Starfish | 0.890±0.074 | 0.881±0.073 | 0.809±0.120 | 0.745±0.067 |
| | Big-FISH | 0.884±0.056 | 0.879±0.060 | 0.797±0.090 | 0.745±0.069 |
| | RS-FISH | 0.923±0.064 | 0.916±0.067 | 0.863±0.110 | 0.783±0.070 |
| | deepBlink | 0.924±0.052 | 0.915±0.056 | 0.863±0.088 | 0.691±0.047 |
| | Spotiflow | **0.941±0.044** | **0.932±0.053** | **0.892±0.077** | 0.825±0.103 |
| | Spotiflow (general) | 0.924±0.054 | 0.915±0.060 | 0.862±0.088 | **0.852±0.090** |
| HybISS | LoG/Starfish | 0.531±0.222 | 0.526±0.221 | 0.390±0.196 | 0.476±0.209 |
| | Big-FISH | 0.644±0.198 | 0.638±0.198 | 0.504±0.204 | 0.566±0.187 |
| | RS-FISH | 0.551±0.241 | 0.543±0.237 | 0.417±0.232 | 0.437±0.194 |
| | deepBlink | 0.738±0.178 | 0.716±0.176 | 0.612±0.196 | 0.553±0.143 |
| | Spotiflow | **0.796±0.146** | **0.792±0.149** | **0.683±0.183** | **0.708±0.145** |
| | Spotiflow (general) | 0.789±0.164 | 0.786±0.166 | 0.678±0.198 | 0.699±0.156 |
| MERFISH | LoG/Starfish | 0.790±0.076 | 0.778±0.077 | 0.659±0.101 | 0.675±0.085 |
| | Big-FISH | 0.222±0.171 | 0.210±0.160 | 0.135±0.113 | 0.154±0.115 |
| | RS-FISH | 0.804±0.081 | 0.791±0.078 | 0.678±0.105 | 0.650±0.063 |
| | deepBlink | 0.834±0.024 | 0.812±0.028 | 0.717±0.035 | 0.615±0.035 |
| | Spotiflow | **0.861±0.030** | **0.847±0.035** | **0.757±0.045** | 0.690±0.048 |
| | Spotiflow (general) | 0.857±0.022 | 0.846±0.027 | 0.751±0.034 | **0.712±0.047** |
| smFISH | LoG/Starfish | 0.821±0.279 | 0.818±0.278 | 0.761±0.284 | 0.700±0.240 |
| | Big-FISH | 0.805±0.228 | 0.794±0.233 | 0.721±0.258 | 0.631±0.205 |
| | RS-FISH | 0.793±0.266 | 0.782±0.268 | 0.720±0.291 | 0.625±0.231 |
| | deepBlink | 0.951±0.069 | 0.949±0.070 | 0.913±0.104 | 0.893±0.141 |
| | Spotiflow | **0.954±0.104** | **0.953±0.104** | **0.925±0.129** | **0.914±0.159** |
| | Spotiflow (general) | 0.947±0.110 | 0.945±0.109 | 0.912±0.133 | 0.892±0.161 |
| Telomeres | LoG/Starfish | 0.700±0.159 | 0.699±0.159 | 0.557±0.165 | 0.605±0.141 |
| | Big-FISH | 0.862±0.043 | 0.861±0.043 | 0.759±0.067 | 0.749±0.036 |
| | RS-FISH | 0.862±0.048 | 0.860±0.048 | 0.760±0.072 | 0.763±0.038 |
| | deepBlink | 0.908±0.027 | 0.906±0.028 | 0.832±0.045 | 0.843±0.031 |
| | Spotiflow | **0.924±0.024** | **0.923±0.025** | **0.860±0.041** | **0.862±0.030** |
| | Spotiflow (general) | 0.913±0.022 | 0.912±0.022 | 0.841±0.036 | 0.843±0.025 |
| Terra | LoG/Starfish | 0.352±0.265 | 0.350±0.264 | 0.248±0.226 | 0.298±0.224 |
| | Big-FISH | 0.541±0.273 | 0.536±0.273 | 0.414±0.246 | 0.440±0.233 |
| | RS-FISH | 0.525±0.209 | 0.520±0.211 | 0.382±0.193 | 0.441±0.195 |
| | deepBlink | 0.717±0.191 | 0.709±0.195 | 0.586±0.191 | 0.609±0.190 |
| | Spotiflow | **0.784±0.099** | **0.778±0.097** | **0.654±0.122** | **0.687±0.090** |
| | Spotiflow (general) | 0.778±0.116 | 0.772±0.114 | 0.648±0.137 | 0.674±0.105 |

**Supplementary Table 4. Performance of different spot methods on a variety of datasets.** For each metric, we show the mean and standard deviation computed across all test images.

**Supplementary Table 5. HybISS (*Mus musculus*) and HCR RNA-FISH (*Platynereis dumerilii*) probe sequences.**

| Dataset | Method | F1 | Accuracy | PLQ |
|---|---|---|---|---|
| Cluster | LoG/starfish | 0.420±0.041 | 0.267±0.033 | 0.261±0.035 |
| | Big-FISH | 0.370±0.087 | 0.231±0.066 | 0.222±0.053 |
| | RS-FISH | 0.767±0.044 | 0.623±0.058 | 0.434±0.040 |
| | deepBlink | 0.850±0.040 | 0.741±0.061 | 0.591±0.040 |
| | Spotiflow | **0.909**±**0.031** | **0.834**±**0.053** | **0.708**±**0.057** |

**Supplementary Table 6. Performance of different spot detection methods on synthetic dense spots clusters.** Metrics were computed on a per-volume basis and were then aggregated.

| Dataset | Method | F1 | Accuracy | PLQ |
|---|---|---|---|---|
| Synthetic (3D) | deepBlink (2D) | 0.102±0.080 | 0.056±0.045 | 0.044±0.032 |
| | Spotiflow (2D) | 0.312±0.087 | 0.188±0.057 | 0.185±0.056 |
| | LoG | 0.608±0.191 | 0.460±0.181 | 0.478±0.151 |
| | Big-FISH | 0.683±0.187 | 0.543±0.188 | 0.531±0.159 |
| | RS-FISH | 0.608±0.176 | 0.457±0.168 | 0.361±0.117 |
| | Spotiflow | **0.882**±**0.110** | **0.805**±**0.165** | **0.708**±**0.131** |

**Supplementary Table 7. Performance of different spot detection methods on a synthetic 3D dataset.** Metrics were computed on a per-volume basis and were then aggregated.

#### Supplementary videos

**Supp. Video 1:** *Data overview.* Visualization of a single channel of a mouse brain embryo acquired with the HybISS protocol highlighting the challenges of processing iST data.

**Supp. Video 2:** *Spotiflow napari plugin.* Demonstration of the Spotiflow napari plugin on a crop of 2D HybISS data as well as a 2D+time live-cell movie containing labelled telomeres.

**Supp. Video 3:** *Stereographic flow.* Animation showcasing the stereographic flow principle.

**Supp. Video 4:** *Live-cell tracking.* Side-to-side comparison of telomeres tracking results obtained from the detections of LoG, deepBlink and Spotiflow on live-cell movies.

**Supp. Video 5:** *3D spot detection on smFISH data.* Side-to-side comparison of *Platynereis dumerilii* smFISH data: raw signal, LoG detections and Spotiflow detections.

**Supp. Video 6:** *3D lipid droplet tracking.* Lipid droplet tracking results obtained from the detections of 3D Spotiflow on a live-cell, label-free volumetric movie.

### 3 Supplementary Notes

#### 3.1 Spotiflow

For simplicity, we focus on the 2D case, but we note that the extension to 3D is direct (see Methods for a brief description of the 3D equations). Given an image and the corresponding spot center annotations we aim to predict two different outputs: i) a *multi-scale probability heatmap*, and ii) an *inverse stereographic vector field* that both encode the location of spots in the image. The network architecture is shown in Supplementary Fig. 1.

##### Multiscale heatmap regression

Given an image $X \in \mathbb{R}^{w\times h}$ and the corresponding spot center annotations $\{p_i\} \subset \mathbb{R}^2$, we first build a probability heatmap $Y \in \mathbb{R}^{w\times h}$ by generating a Gaussian distribution of variance $\sigma$ centered at every spot so that the probability map exponentially decays around the annotate centre.

$$Y(x) = \max_{p\in\{p_i\}} \exp\left(-\frac{\|p-x\|_2^2}{2\sigma^2}\right) \in [0,1] \tag{1}$$

Note that instead of summing the individual Gaussian distributions, we take the maximum value at each pixel to create sharp boundaries between spots. We then train a multiscale convolutional neural network on image-heatmap pairs $(X, Y)$ which aims to regress the Gaussian heatmap $Y$ given the image $X$. While any multiscale architecture can be used as a backbone, we use a UNet [12].

In order to help convergence, avoid vanishing gradients, and effectively guide the solution of higher resolution maps, we add small convolutional heads at different resolution levels (the decoder feature maps in the case of the UNet) and regress the Gaussian heatmap $Y^{(l)}$ at different levels $l = 0 \dots L-1$ (corresponding to a resolution reduction by $\frac{1}{2^l}$) using the following pixel-wise multiscale loss w.r.t. to the ground truth heatmaps $\hat{Y}^{(l)}$:

$$\mathcal{L}_{heat}(Y,\hat{Y}) = -\left(1+\lambda\mathbb{1}_{\hat{Y}^{(0)}>\epsilon}\right)\sum_{l=0}^{L-1}\frac{1}{2^l}\left[\hat{Y}^{(l)}\log\left(Y^{(l)}\right)+\left(1-\hat{Y}^{(l)}\right)\log\left(1-Y^{(l)}\right)\right] \tag{2}$$

Here $\lambda$ is used to control the weight given to positive samples (*i.e.* spots, pixels whose value in the heatmap is greater than a threshold value $\epsilon$) in the loss function: a higher value of $\lambda$ will penalize missing spots, which will bring up the number of detections but, at the same time, will increase the number of false positives. On the other hand, smaller values of $\lambda$ may cause a higher number of false negatives. The prefactor $\frac{1}{2^l}$ is used to downweight lower resolution terms to avoid them dominating the optimization. During inference, a simple local maxima detector on the full-resolution heatmap suffices to extract the spot centroids whose probability is higher than a certain threshold. This probability threshold is automatically optimized on the validation dataset during training.

##### Stereographic flow

For each pixel $x = (i,j) \in \mathbb{R}^2$ of the image $X$, we first define a *local vector field* $v_{ij} = (v_x, v_y) \in \mathbb{R}^2$ given by the vector from the pixel to the nearest ground truth spot (Supplementary Fig. 2a). As a result, $v_{ij} = (0,0)$ if and only if there is a spot centered at the pixel $(i,j)$. Note that the vector field defined this way would have arbitrary magnitude (*i.e.* $||v_{ij}||_2 \to \infty$) in the situation of far away spots. This can cause numerical unstability if this vector field is to be regressed, which is exactly the task at our hands. To fix this problem, we make use of a scaled *inverse stereographic projection* $f : (v_x, v_y) \in \mathbb{R}^2 \to (v'_x, v'_y, v'_z) \in S^2$ which satisfies

$$v_x = \frac{v'_x}{1+v'_z}, \quad v_y = \frac{v'_y}{1+v'_z}, \quad 1 = {v'_x}^2 + {v'_y}^2 + {v'_z}^2$$

and maps points on the sphere $S^2 \setminus \{(0,0,-1)\}$ to the Euclidean plane $\mathbb{R}^2$. Effectively, we represent the vector field as a point on the unit 3D sphere $S^2$ (this generalizes to arbitrary dimensions). In particular, $f^{-1}$ maps the zero vector $(0,0)$ to the north pole $(0,0,1)$ and all vectors with infinite length ("points at infinity") to the south pole $(0,0,-1)$. Additionally we introduce a fixed length scale $s$ and scale the initial vector field to $\left(\frac{v_x}{s}, \frac{v_y}{s}\right)$, which yields the embedding:

$$v'_x = \frac{2sv_x}{r^2+s^2}, \quad v'_y = \frac{2sv_y}{r^2+s^2}, \quad v'_z = -\frac{r^2-s^2}{r^2+s^2} \quad \text{with } r^2 = v_x^2 + v_y^2$$

(Supplementary Fig. 2b). The embedding can be inverted via the corresponding scaled stereographic projection $f$:

$$v_x = \frac{sv'_x}{1+v'_z}, \quad v_y = \frac{sv'_y}{1+v'_z}$$

Note that the $v'_z$ coordinate resembles a shifted gaussian distribution centered at the spot:

$$v'_z = -\frac{r^2 - s^2}{r^2 + s^2} \approx 2e^{-r^2/s^2} - 1$$

with standard deviation $\sigma = \frac{s}{\sqrt{2}}$.

During training, we then not only regress the Gaussian heatmaps, but also the stereographic flow computed using an extra convolutional head at the highest resolution. We optimize a full resolution pixel-wise weighted L1 loss $\mathcal{L}_{flow}$ between the ground truth stereographic flow $V' = \{v'_{ij}\}$ and the predicted field $\hat{V}' = \{\hat{v'}_{ij}\}$:

$$\mathcal{L}_{flow}(V', \hat{V}') = \sum_{i,j} (1 + \mathbb{1}_{\|v'_{ij}\|_2 < \epsilon}\lambda) \|v'_{ij} - \hat{v'}_{ij}\|_1 \tag{3}$$

where $v'_{ij} = (v'_{x,ij}, v'_{y,ij}, v'_{z,ij}) \in S^2$ and $\hat{v'} = (\hat{v}'_{ij,x}, \hat{v}'_{ij,y}, \hat{z}'_{ij}) \in S^2$. Note that, by construction, $\min_k v_k \geq -1$ and $\max_k v_k \leq 1$, inducing numerical stability. The hyperparameter $\lambda$ again controls the weight given to spot locations. We note that in our implementation, we do not use $||v'||_2 < \epsilon'$ but equivalently use the ground truth heatmap to reuse computations. The overall loss function optimized by the network is

$$\mathcal{L} = \mathcal{L}_{heat} + \mathcal{L}_{flow} \tag{4}$$

The stereographic field not only helps maintain numerical stability during training, but also achieves subpixel resolution. Such precision can be obtained by projecting the predicted stereographic flow $\hat{v'}$ to $\hat{v} = f(\hat{v'}) \in \mathbb{R}^2$ and adding the offset $\hat{v}_{p_x,p_y}$ to the spot center predictions $(p_x, p_y)$ obtained using the heatmap $Y$.

##### 3.2 Evaluation Metrics

For each image, let $\{p_i\}, p_i \in \mathbb{R}^2$ be the set of ground truth spot coordinates and $\{\hat{p}_j\}, \hat{p}_j \in \mathbb{R}^2$ the set of predicted spot coordinates.

**Detection metrics** To compute overall detection metrics for each image, we first uniquely match ground truth $\{p_i\}$ and predicted spots $\{\hat{p}_j\}$ according to their spatial proximity via hungarian matching [13]. We then define a spatial cutoff $c \in \mathbb{R}$ and count a matched pair $(p, \hat{p})$ as *true positive* (TP) if their Euclidean distance $d(p, \hat{p}) = \|p - \hat{p}\|^2$ is not larger than $c$, a predicted spot $\hat{p}$ as *false positive* (FP) if there was no matched ground truth spot, and a ground truth spot $p$ as *false negative* (FN) if there was no matched predicted spot. We then define for each image the following metrics:

$$F_{1[c]} = \frac{|TP|}{|TP| + \frac{1}{2}(|FP| + |FN|)} \tag{5}$$

$$AP_{[c]} = \frac{|TP|}{|TP| + |FP| + |FN|} \tag{6}$$

(7)

Furthermore, in order to report a metric which takes into account different spatial cutoffs $c$, we also report the $F_1 AuC$ [4]:

$$F_1 AuC_{[c_L;c_H]} = \frac{1}{c_H - c_L} \sum_{k=L}^{H-1} \frac{F_{1[c_k]} + F_{1[c_{k+1}]}}{2} \Delta; \quad \text{with} \quad c_L < c_H \tag{8}$$

where $\Delta$ is the constant difference between two successive cutoffs $c_k$ and $c_{k+1}$ ($\Delta = c_{k+1} - c_k > 0$). Note that the numerator is a (rescaled) trapezoidal approximation to the integral of the curve generated by the $F_1$ metric at different cutoffs $c_k$. We set $\Delta = 0.25$ throughout the experiments. The denominator ensures $F_1 AuC \in [0, 1]$.

**Panoptic localization quality (PLQ)** To additionally incorporate the spatial localization accuracy of the predicted spots for each image, we follow [14] and define the average *localization accuracy* $LA$

$$LA_{[c]} = \frac{1}{|TP|} \sum_{(p,\hat{p}) \in TP} 1 - \frac{\max(d(p,\hat{p}), c)}{c} \tag{9}$$

and the *panoptic localization quality* $PLQ$ as

$$PLQ_{[c]} = LA_{[c]} \cdot F_{1\,[c]}. \tag{10}$$

Note that the localization accuracy is equal to 1, if all correctly matched detections perfectly colocalize with the ground truth spots, and it equals 0 if no detections are matched to the ground truth with the given cutoff $c$.

#### 3.3 Spot detection benchmarking details

For all compared methods we normalized the intensities of input images via percentile-based min-max normalization ($p_{min} = 3, p_{max} = 99.8$) before feeding them to the methods. To ensure a fair comparison, we optimize their parameters on each respective dataset in the following way:

##### Starfish

We find the optimal intensity threshold in Starfish in a two-step process:

- Coarse-grained optimization: we test 11 values in the interval $[0.1, 1]$, which are evenly spaced in a logarithmic scale, and retrieve an interval $[t_0, t_1]$ containing the best-found coarse threshold $t_c$.
- Fine-grained optimization: we test 11 values in the interval $[t_0, t_1]$, also evenly spaced in a logarithmic scale, and retrieve the final threshold $t$.

##### Big-FISH

We find the optimal $\sigma$, which is the variance of the LoG filter, in a coarse-to-fine approach similarly to the one just described:

- Coarse-grained optimization: we test 11 values for $\sigma$ in the interval $[1, 10]$, which are evenly spaced in a logarithmic scale, and retrieve an interval $[\sigma_0, \sigma_1]$ containing the best-found coarse variance $\sigma_c$.
- Fine-grained optimization: we test 11 values in the interval $[\sigma_0, \sigma_1]$, also evenly spaced in a logarithmic scale, and retrieve the final threshold $\sigma$.

. The intensity threshold is automatically found by the algorithm using an elbow rule heuristic.

##### deepBlink

We used default values as described in the original paper and as output by the command *deepblink config*.

##### RS-FISH

Grid search was run on the original ImageJ RS-FISH implementation jointly for all parameters but the intensity threshold, which was optimized in an offline fashion to reduce computational load. During the grid search, the intensity threshold was set to a conservative value to retrieve as many spots as possible, which were then filtered by intensity in the last parameter optimization step.

Taking the intensity threshold values into account, a total of 147420 parameter configurations were tested.

- Outlier removal = "RANSAC"
- $\sigma$ of DoG $\in [1, 2.5]$, uniform spacing of 0.25
- Threshold of DoG $\in [0.001, 0.13]$, multiplicative spacing of 1.50
- Support radius $\in \{2, 3, 4\}$
- Inlier ratio $\in \{0, 0.1, 0.2, 0.3\}$
- Max error $\in [0.1, 2.6]$, uniform spacing of 0.5
- Background substraction method = "Mean"
- Intensity threshold: 30 values uniformly spaced in the interval $[I_{min}, I_{max}]$, where $I_{min}, I_{max}$ are, respectively, the minimum and maximum intensities of spot candidates across a whole dataset.

###### Spotiflow

No grid search has been performed on Spotiflow. The default values used are

- Batch size = 4
- Learning rate = 3e-4
- Patch size = 512
- Resolution levels = 4
- Number of convolutions per level = 3
- Backbone architecture = UNet
- Loss weight for positive class = 10
- Number of epochs = 200
- Number of initial feature maps = 32
- Feature map increase factor = 2
- Augmentations: flip-rotations, rotations, Gaussian noise addition, intensity shifts
- $\sigma$ for Gaussian heatmap generation = 1
- Grid factor $g = 2$ (3D models only)

#### 3.4 Scalability experiments

In the 2D case, the images used were obtained by consecutively expanding a center crop from a full HybISS cycle. For the 3D assessment, the volumes used were obtained by concatenating the same synthetic volume from the dataset *synthetic-3D* along the three axes.

Given the dependency of intensity-based methods on the number of spots, their parameters were set such that the number of detections are in the same order of magnitude across all methods. Results were obtained using the Python-based profiling tool Scalene for all methods but RS-FISH, whose results were obtained Unix's *time* module. Intensity-based methods were run on an AMD Ryzen Threadripper PRO 5965WX 24-Cores CPU with 256GB of memory.

Learning methods (deepBlink and Spotiflow) were run on an NVIDIA GeForce RTX 4090 GPU (24GB). Intensity-based methods tend to scale poorly with the image size compared to Spotiflow. This is particularly important as the resolution of FISH-based spatial transcriptomics is very high to enable single-cell analyses.

#### 3.5 Training a Spotiflow model

Before deciding to train a Spotiflow model, we recommend testing out the pre-trained models and assess whether the accuracy is sufficient (*cf.* Supp. Fig. 20). For both training and fine-tuning (which requires less amount of annotations, *cf.* Supp. Fig. 12) the spot annotations need to be in a specific format in order to use Spotiflow's CLI directly:

```
data folder
├── train
│   ├── img_001.csv
│   ├── img_001.tif
│   ├── ...
│   ├── img_064.csv
│   └── img_064.tif
├── val
│   ├── img_065.csv
│   ├── img_065.tif
│   └── ...
└── test
    ├── ...
    ├── img_100.csv
    └── img_100.tif
```

Note that the filenames can be arbitrary, and the only requirement is that the stem of the filenames are the same for the corresponding image-annotation pairs. The annotations should be in CSV format containing two columns (for 2D) or three (for 3D), with the expected header names being ‘axis-0‘, ‘axis-1‘, ‘axis-2‘ (the latter only in 3D). One row in the CSV should correspond to the annotation centre for in pixel space. Below is the first three rows for an example file of 2D annotations:

```
axis-0,axis-1
324.4,123.2
23.43,412.8
...
```

After having the annotations in the appropriate format, a Spotiflow model with the default hyperparameters can be trained by running the following from the command line:

```
$ spotiflow-train /path/to/data_folder -o /my/trained/model
```

which will save the trained model to the folder */my/trained/model*. If instead one wants to fine-tune on an existing model, adding a new parameter to the CLI suffices:

```
$ spotiflow-train /path/to/data_folder -o /my/finetuned/model --finetune-from general
```

To see the full set of parameters, one can use the help functionality of the command:

```
$ spotiflow-train -h
```

We note that this works as for version 0.5.0, and we recommend checking the documentation of the software for the latest release (URL), which includes more extensive information on the training procedure.
